# Evolutionary Design of Membrane-Lytic Antimicrobial Peptides with Mixture of Experts

**DOI:** 10.64898/2026.07.29.741660

**Authors:** Ruihan Dong, Chen Song

**Affiliations:** Center for Quantitative Biology, Academy for Advanced Interdisciplinary Studies, Peking University, Beijing 100871, China; Peking-Tsinghua Center for Life Sciences, Academy for Advanced Interdisciplinary Studies, Peking University, Beijing 100871, China; Peking University–Tsinghua University–National Institute of Biological Sciences Joint Graduate Program, Academy for Advanced Interdisciplinary Studies, Peking University, Beijing 100871, China

**Keywords:** Antimicrobial peptide, Membrane-lytic, Mixture of experts, Sequence design, Generative adversarial imitation learning

## Abstract

Antimicrobial peptides (AMPs) hold great promise in combating drug-resistant pathogens, yet current design models do not explicitly account for their primary mode of action, membrane disruption. To address this gap, we propose AMPainterV2, a model that designs membrane-lytic AMPs using generative adversarial imitation learning. AM-PainterV2 integrates a Mixture-of-Experts (MoE) policy network to support insertion, deletion, and mutation operations, thereby expanding the evolutionary design space. In vitro experiments demonstrate that all ten top-ranked peptides evolved from random sequences are active membrane-lytic AMPs, with the best candidate achieving a mean minimal inhibitory concentration (MIC) of 1.5 µM and superior membrane-lytic activity compared to polymyxin B. Particularly, a deletion-only version of AMPainterV2 enables membrane-lytic AMP miniaturization, yielding six miniaturized peptides with comparable or improved activity and an average 36% length reduction. Collectively, AMPainterV2 advances mechanism-driven AMP design and offers a valuable tool for cost-effective AMP development.

## Introduction

Antimicrobial peptides (AMPs) are emerging therapeutic agents to combat multidrug-resistant bacteria. These short peptides exert diverse mechanisms of action, including disruption of membranes, interference with intracellular targets (e.g., nucleic acids and proteins), and modulation of host immune responses. ^1,2^ Among these, membrane disruption is the most recognized mechanism. Membrane-lytic AMPs are considered less likely to induce resistance due to the conserved nature of lipid bilayers. ^3^ Recent progress in artificial intelligence has greatly accelerated the discovery of novel AMPs through both sequence mining and generation, ^4^ with many candidates demonstrating membrane-lytic activity in vitro. ^5–8^ However, such mechanistic characterization is often overlooked during computational design. Our group previously developed a support vector machine (SVM) predictor for screening membrane-lytic AMPs from proteomes. ^9^ Yet, to our knowledge, no existing AMP generative or optimization model has been explicitly designed to target this functional class.

AMP optimization methods can be classified into two categories based on the definition of the optimization space. Continuous approaches embed peptide sequences into a latent space (typically via variational autoencoders), and perturb the embedding through sampling strategies such as Bayesian optimization, ^10^ conditional sampling, ^11^ or zero-order optimization. ^12^ Discrete approaches iteratively edit sequences guided by a fitness function, usually using mutation and crossover operations inspired by genetic algorithms. ^13,14^ EvoGradient, for in-stance, leverages a neural network predictor to introduce mutations as gradient-based directed evolution. ^15^ Although such discrete methods can provide traceable evolutionary paths and are referred to as *evolutionary design*, they are constrained by the low efficiency of single-point mutations and the absence of variable-length operations (i.e., insertion and deletion), which restricts the design space.

The Mixture-of-Experts (MoE) framework was originally proposed to scale up neural net-work capacity by replacing dense layers with sparse modules. ^16^ An MoE architecture comprises a router and several expert subnetworks, where the router dynamically selects which expert to activate, and each expert autonomously specializes in distinct aspects of the data without explicit prior assignment. In our previous AMP design model, AMPainter, ^17^ we decomposed mutation into site selection and residue replacement, enabling effective AMP evolution from diverse initial sequences. To extend this approach to support multiple sequence edit operations, the network must jointly specify both the operation and the corresponding residue. This requirement aligns naturally with the MoE architecture, as different experts can be dedicated to different edit operations.

Therefore, we propose AMPainterV2, a membrane-lytic AMP design model that integrates an MoE policy network within a generative adversarial imitation learning (GAIL) frame-work. Rather than using the MoE framework for model scaling, we adopt it to handle insertion, deletion, and mutation operations, while GAIL allows it to learn directly from available membrane-lytic AMPs as demonstration data. Even without explicit guidance from an antimicrobial scorer, AMPainterV2 can significantly improve the activity of both known AMPs and random sequences, outperforming other AMP optimization methods. In vitro experiments demonstrate a success rate of 100% for ten membrane-lytic AMPs evolved from random inactive sequences. Eight of these AMPs showed broad-spectrum activity against four bacterial species. We also provide a deletion-only version of the AMPainterV2 model, which effectively miniaturizes the AMPs while preserving their membrane-lytic activity. The best miniaturized AMP reduces its length by 45% while increasing its antimicrobial activity by approximately 40 times, with a mean MIC of 2.5 µM. In short, AMPainterV2 demonstrates a mechanism-driven AMP design approach and expands the discrete design space.

## Results

### Training AMPainterV2 with GAIL

AMPainterV2 builds on the approach of our AMP design framework AMPainter. ^17^ We adopted the same evolutionary design mode by decoupling the steps of assigning the edit site and residue type. Accordingly, AMPainterV2 comprises a policy network in conjunction with a protein language model that has been fine-tuned on AMP sequences (Fig. 1a). Compared to AM-Painter, AMPainterV2 features two major advancements: (i) an expanded MoE policy network that supports insertion, deletion, and mutation as distinct sequence edit operations; and (ii) the integration of GAIL to direct the design process specifically toward membrane-lytic AMPs.

**Figure 1:**
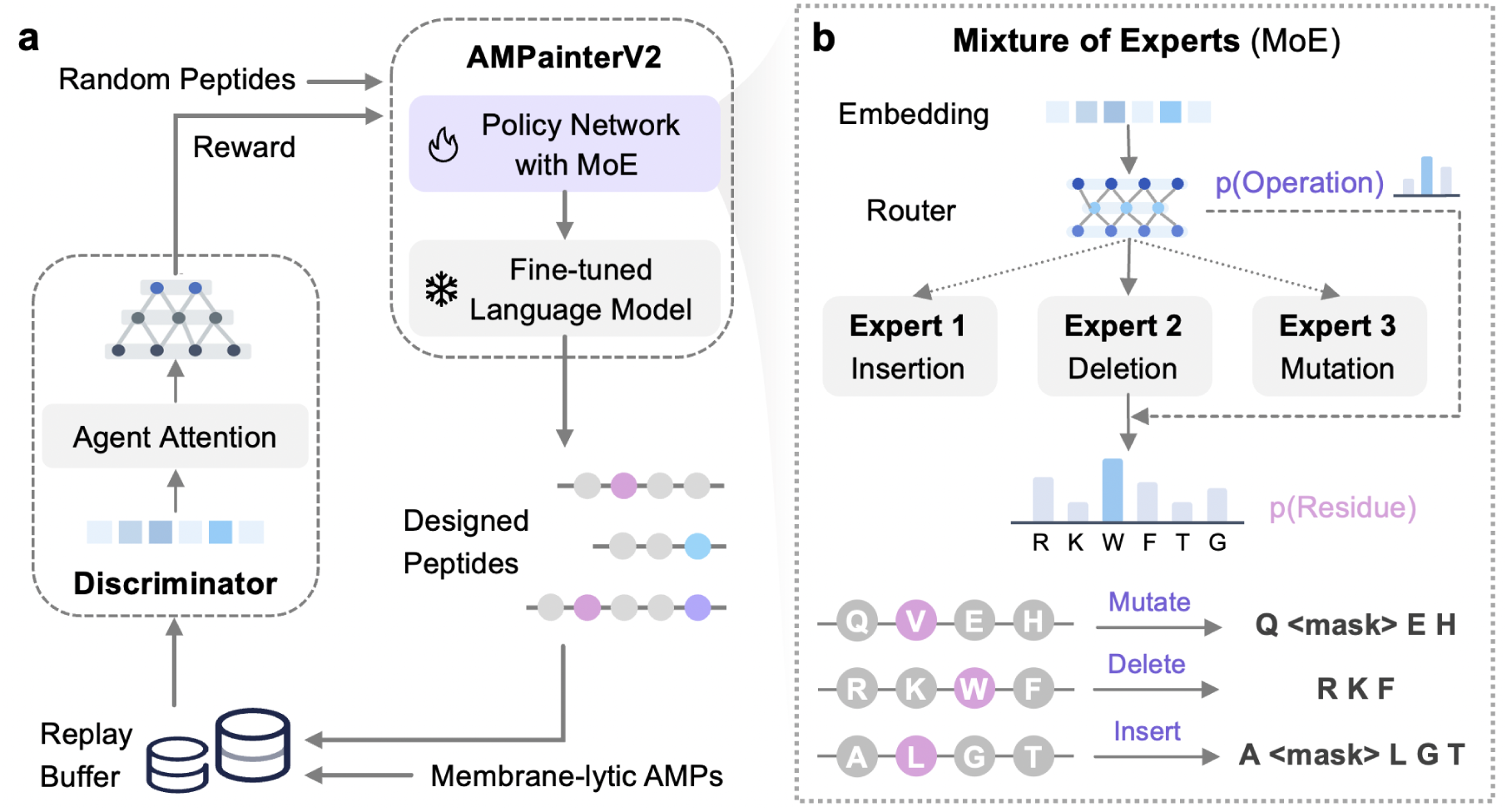
AMPainterV2 framework. **a**. AMPainterV2 was trained with the generative adversarial imitation learning (GAIL), using membrane-lytic AMPs as demonstration data. **b**. The Mixture-of-Experts (MoE) policy network to process mutation, insertion, and deletion operations on sequences.

We incorporated a MoE policy network to handle three sequence edit operations. As shown in Fig. 1b, a router is responsible for producing the probability distribution over three operations. The operation with the highest probability is subsequently selected, thereby activating the corresponding expert subnetwork (Eq. (1)). The activated expert then produces the weight for each residue site, which is used to form a probability distribution over sites (Eq. (2)). The AMPainterV2 agent determines the edit site of the input sequence by sampling from this distribution. A masking token (*<mask>*) will be inserted prior to the site for the insertion operation or will replace the assigned site for mutation. Then the fine-tuned language model decodes the masking tokens with new residues. Conversely, when deletion is chosen, the assigned site will be deleted directly.

We adopted a GAIL framework ^18^ to train the AMPainterV2 for designing membrane-lytic AMPs. Rather than defining an explicit reward function, we employed a curated set of 481 experimentally verified membrane-lytic AMPs from our MemAMPdb ^19^ as demonstration data. In the training loop shown in Fig. 1a, we collected the designed peptides after they were edited by the agent and stored them in a replay buffer. Then we scored these peptides using a discriminator module as reward signals to update the policy network. The demonstration data were labeled as positive, whereas the designed peptides were treated as negative. Therefore, the discriminator output represents the probability that a sequence belongs to the membrane-lytic AMP. As an adversarial learning process, we alternately updated the discriminator and the policy network until the discriminator could no longer distinguish the designed peptides from the demonstration data. The replay buffer was cleared at the end of each episode.

We trained the model for a total of 40 episodes, during which the perplexity gradually decreased, indicating the improved quality of the designed peptides (Fig. S1a). The Boman index of the sequences also decreased, indicating a better membrane-interacting potential (Fig. S1b). The average antimicrobial activity of designed sequences (HyperAMP ^17^ score) increased from 0.031 (initial random sequences) to a maximum score of 0.442 (Fig. S1c). Therefore, we selected this checkpoint at episode 30 as the final AMPainterV2 model.

### Preferences of Sequence Edit Operations in MoE

We leverage the sparse expert subnetworks in MoE to designate three kinds of sequence edit operations instead of using MoE to scale up the model. To maintain load balance within the MoE and prevent consecutively activating a single expert, we added Gaussian noise to the router output (Eq. (2)). This ensured that the fraction of edit operations was generally evenly distributed during the training episodes (Fig. 2a). “No operation” accounted for one quarter because we set the maximum edit distance of each sequence to one third of the initial sequence length, following the setting of deepAMP. ^20^ If we discarded the noise, the HyperAMP score rose more slowly and took 40 episodes to peak at 0.444 (Fig. S2a), which is similar to the maximum score of AMPainterV2. Meanwhile, the mutation operation dominated the output of MoE policy without noise, accompanied by a small portion of insertions (Fig. S2b). This suggests that the MoE policy without noise still preferred length-preserving edits, which limits the design space.

**Figure 2:**
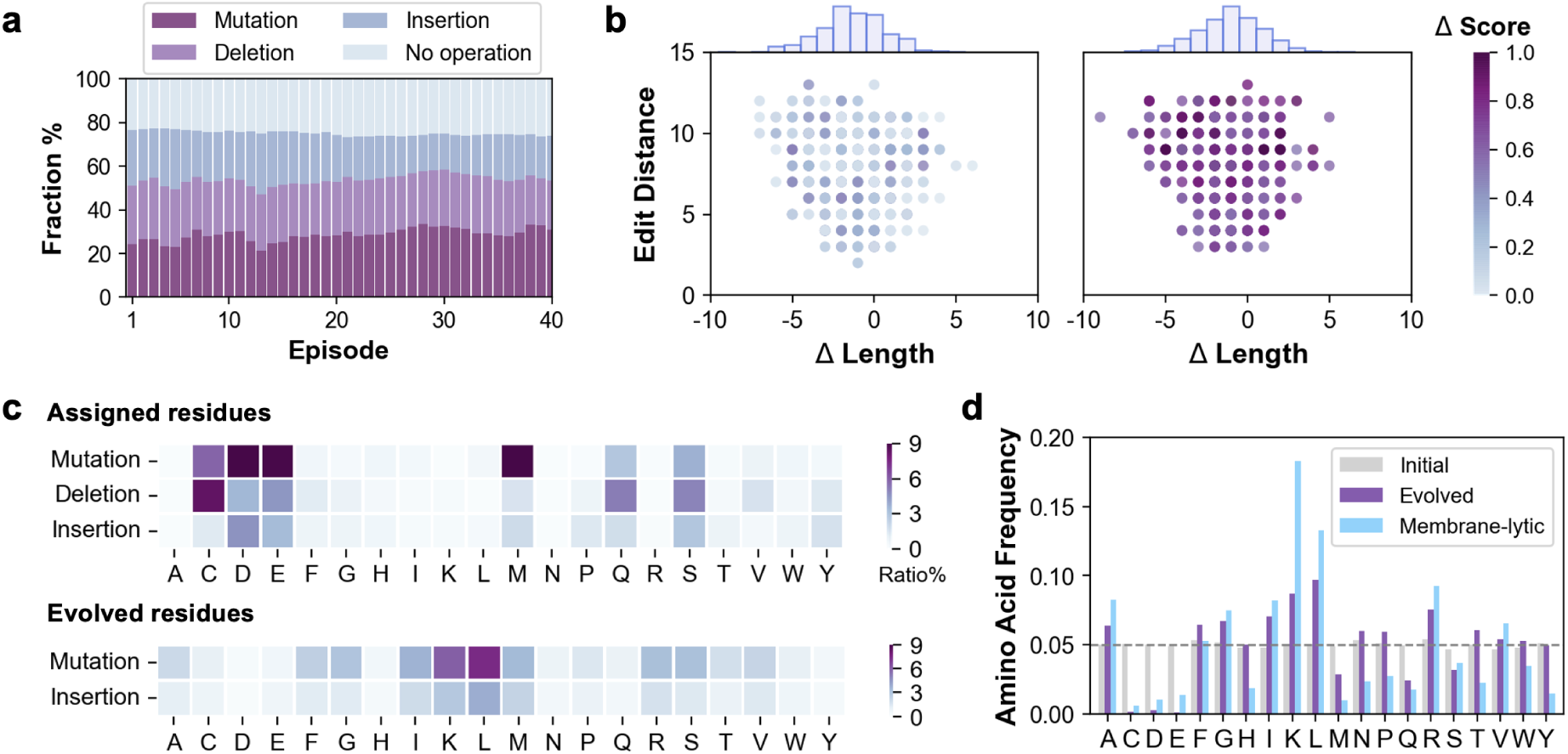
Analysis of the sequence edit operations in AMPainterV2. **a**. Fraction of each edit operation in MoE during 40-episode training. **b**. The distribution of edit distances, length differences, and increasing antimicrobial scores (predicted by HyperAMP) of evolved sequences at episode 30 (Δ Score < 0.5 or ≥ 0.5). **c**. Assigned or evolved amino acid preference of different edit operations. **d**. Amino acid frequency of the initial random sequences, evolved sequences at episode 30, and the membrane-lytic AMPs.

We collected the evolved sequences of each initial random sequence at episode 30 of AMPainterV2 for analysis. The Levenshtein edit distances of these sequences ranged from 2 to 13. Fig. 2b separated the evolved sequences with increasing HyperAMP scores (Δ*Score* < 0.5 or else) for clearer visualization. 82.7% of evolved sequences have Δ*Length ≠* 0 and 42.8% of them are associated with Δ*Score* ≥ 0.5, suggesting that insertion and deletion operations have played roles in enhancing antimicrobial activity. Longer sequences were more likely to be deleted (Fig. S3a), as the lengths of membrane-lytic AMPs tended to be between 10 and 25 (Fig. S3b).

We further conducted an in-depth analysis of amino acid preferences when the MoE policy assigned operations and residues. Fig. 2c showed that the MoE tended to mutate cysteine (C), aspartic acid (D), glutamic acid (E), and methionine (M), while frequently deleting C along with some glutamine (Q) and serine (S). This trend was consistent with the amino acid composition of the membrane-lytic AMPs used as demonstration data in GAIL training. Indeed, Fig. 2d showed that these membrane-lytic AMPs contain low proportions of C, D, E, and M. In contrast, the MoE policy without noise preferred to delete or mutate a fixed set of amino acids (Fig. S2c). Evolved residues were determined by the fine-tuned language model in AMPainterV2, and their preferences for positively charged lysine (K) and hydrophobic leucine (L) matched the trend observed in membrane-lytic AMPs (Fig. 2c, d). Together, these results demonstrated that different sequence edit operations guided the initial random sequences toward membrane-lytic AMPs as the policy was trained.

### AMPainterV2 Surpasses Other Methods

We compared the AMPainterV2 model with other AMP optimization models on two design tasks, using either AMPs or random sequences as inputs (Fig. 3a). The goal is to evolve these sequences into membrane-lytic AMPs with high antimicrobial activity. For the AMP dataset, we collected 144 new AMPs with unknown mechanisms from recently published articles to avoid overlap with the training data of all models. The random sequence dataset was generated with equal frequencies of 20 canonical amino acids. We fed each sequence into each model, ran ten parallel repeats, and ranked the optimized sequences by their HyperAMP scores. We analyzed the top candidates in terms of antimicrobial activity, toxicity, membrane-lytic mechanism, and perplexity.

**Figure 3:**
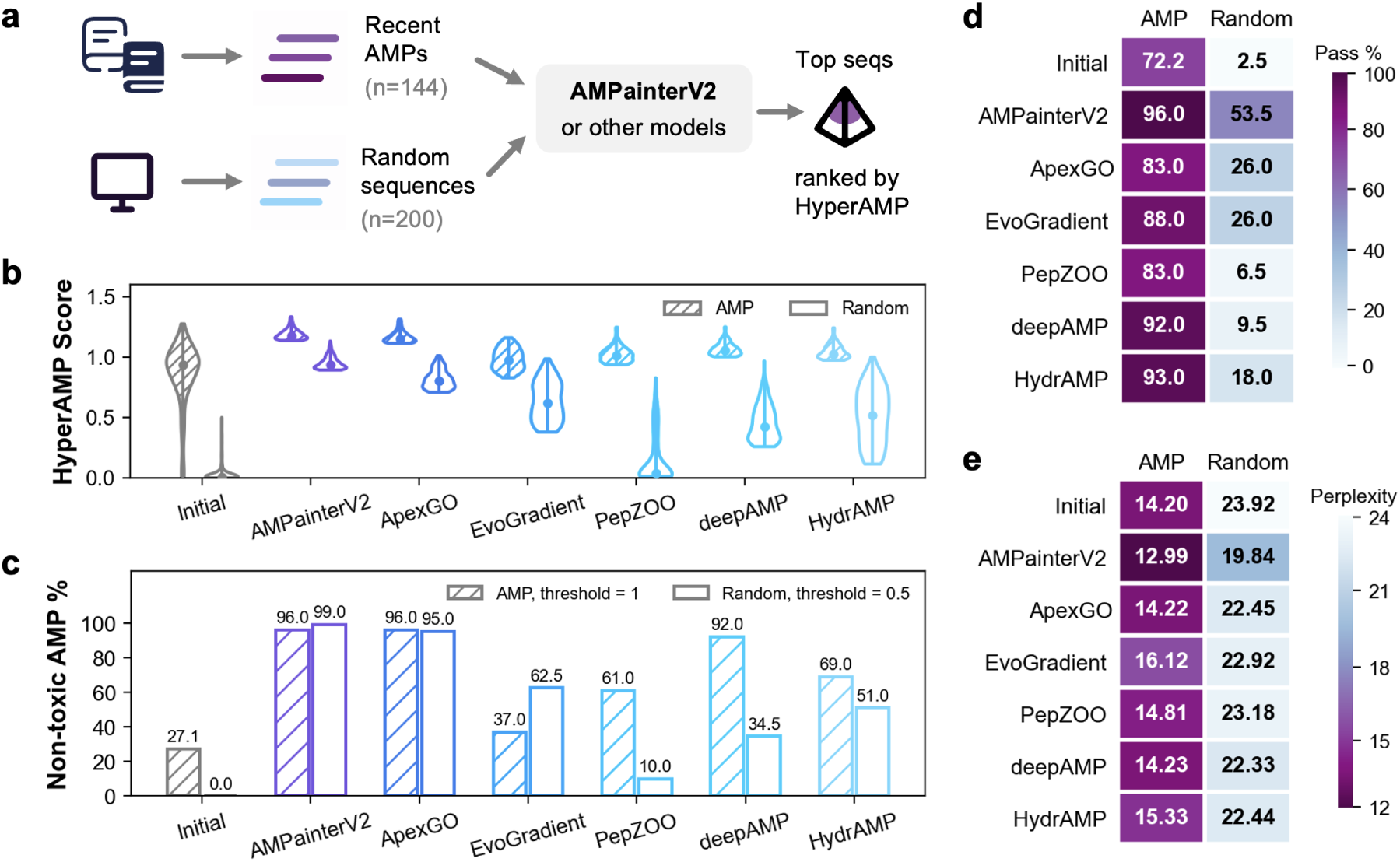
Comparison of AMPainterV2 and other models. **a**. Workflow of comparison. A dataset of AMPs collected from recent articles and a dataset of random sequences were used as inputs to AMPainterV2 and four other AMP optimization models. The top 100 or 200 evolved sequences with the highest HyperAMP scores were analyzed. **b**. HyperAMP scores of initial and evolved sequences. **c**. Fraction of non-toxic AMPs in initial and evolved sequences. Toxicity was evaluated by BioToxinPept. The threshold of HyperAMP score was 1 for sequences evolved from AMPs and 0.5 for those evolved from random sequences, respectively. **d**. Ratio of passing the SVM predictor for membrane-lytic AMPs. **e**. ESM Perplexity of the initial and evolved sequences.

We chose five AMP optimization models for comparison, one of which is a discrete method that involves only the mutation operation (EvoGradient ^15^), while the others are VAE-based continuous methods (ApexGO, ^10^ PepZOO, ^12^ deepAMP, ^20^ and HydrAMP ^11^). All tested models were able to improve the activity of known AMPs, with AMPainterV2 achieving the highest scores, as evaluated by HyperAMP (Fig. 3b). However, when starting from random sequences, only AMPainterV2 could raise the scores to a comparable level, while the other models produced modest improvements. Using another antimicrobial activity predictor also yields similar results (Fig. S4a). This is probably because AMPainterV2 was trained with random sequences as initial inputs and learned the strategy of evolving them into AMPs, while the other AMP optimization models were trained on AMP sequences. Notably, AMPainterV2 also obtained the highest ratio of non-toxic AMPs, reaching 96% and 99% on the two tasks as evaluated by BioToxinPept ^6^ (Fig. 3c), and 58% and 67% by ToxinPred 3.0^21^ (Fig. S4b). In fact, our demonstration data contain fewer toxic AMPs than the training sets of other methods (Table S1). Only ApexGO achieved comparable ratios, as it was pretrained on universal peptides and subsequently fine-tuned on random variants of template AMPs.

Unlike other methods, AMPainterV2 is mechanism-driven in its design objective. We used the SVM predictor developed by our group ^9^ to assess the membrane-lytic mechanism of the evolved sequences (Fig. 3d). 72.2% of the initial AMPs were predicted to inhibit bacteria via membrane disruption. AMPainterV2 increased this ratio to 96.0%, slightly higher than that of other methods. Even though other AMP optimization methods did not take the mechanism into account, they still raised the ratio to above 80% alongside activity improvements. However, AMPainterV2 significantly boosted the SVM pass ratio of random sequences from 2.5% to 53.5%, while other methods showed no comparable improvement. Meanwhile, the evolved sequences of AMPainterV2 were more similar to natural peptides, as indicated by lower evolutionary scale modeling (ESM) perplexity scores (Fig. 3e). Our method also outperformed others in terms of both sequence diversity and similarity (Fig. S4c).

In addition, we assessed a variant of AMPainterV2 trained using AMPs without specifically assigned mechanisms (non-specific AMPs) as the demonstration data, in place of membrane-lytic AMPs. Comparative analysis in Table S2 revealed that while both models produced comparable proportions of non-toxic AMPs, the variant trained on non-specific AMPs exhibited a substantially lower SVM pass rate than the original model. This finding highlights the importance of demonstration data in mechanism-driven design. In summary, AMPainterV2 enables the evolution of both AMPs with unknown mechanisms and random sequences into non-toxic, highly active membrane-lytic AMPs, demonstrating its potential in antimicrobial drug development.

### Membrane-Lytic AMPs Evolved by AMPainterV2

We selected ten top-ranked peptides evolved from random sequences into the experimental stage (named V2-R01 to V2-R10, Fig. 4a). BLAST-p searches against the DRAMP ^22^ and AMP-Sphere ^23^ databases revealed no matches for them, with only a distant hit for V2-R09 (E-value = 3.2), confirming that these are de novo peptides. All ten peptides exhibited low Boman index values, indicating a high potential for interacting with membranes (Table S3).

**Figure 4:**
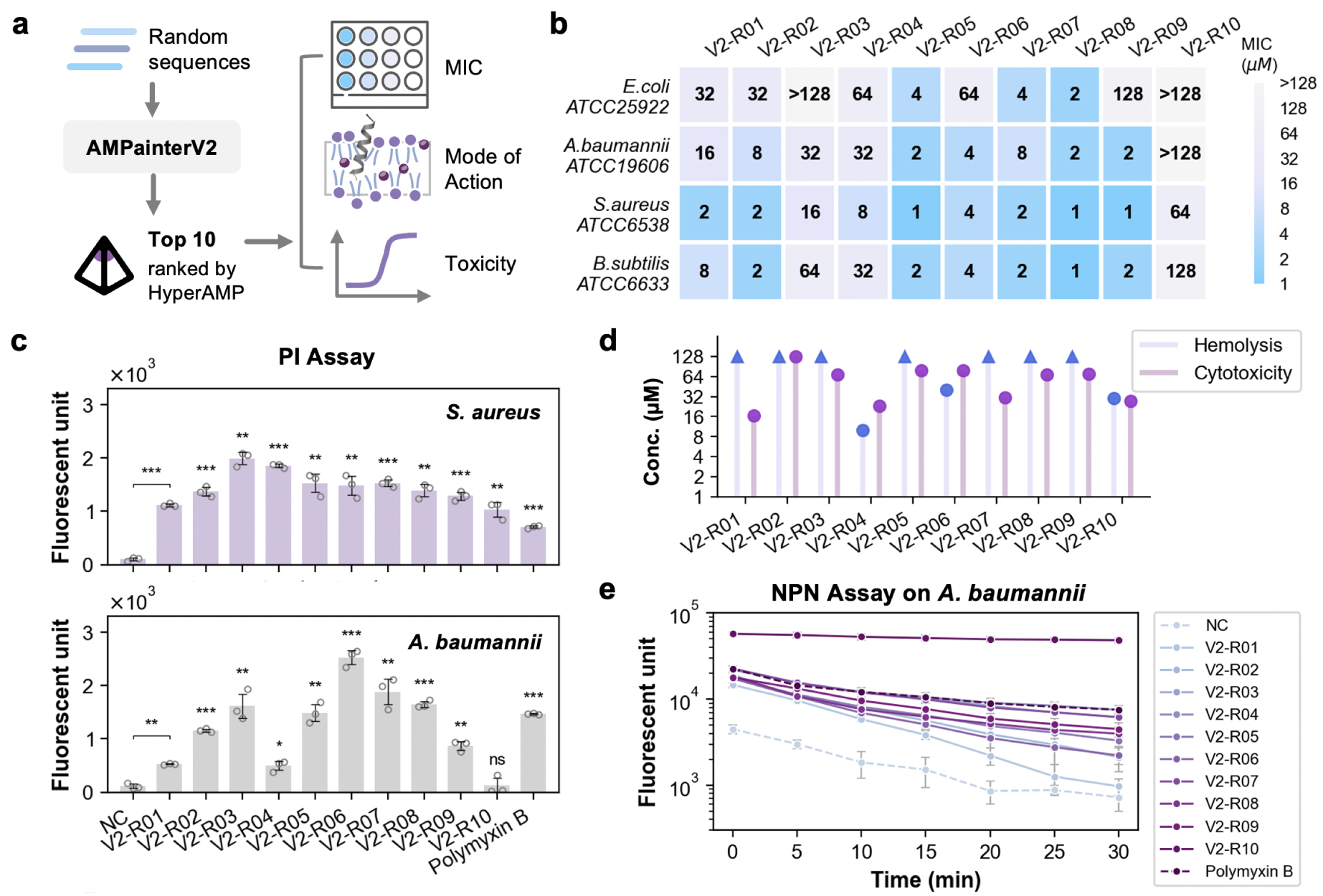
Experimental validation of top ten designed peptides, as ranked by HyperAMP. **a**. The top ten sequences evolved from random sequences using AMPainterV2 were selected for minimal inhibitory concentration (MIC), mode of action, and toxicity tests. **b**. MIC measurements of ten peptides against *Escherichia coli* ATCC25922, *Acinetobacter baumannii* ATCC19606, *Staphylococcus aureus* ATCC6538, and *Bacillus subtilis* ATCC6633. **c**. Cytoplasmic membrane disruption assay of ten peptides on *S.aureus* ATCC6538 and *A.baumannii* ATCC19606 using propidium iodide (PI). Unpaired t-tests were used for comparison with the negative control (NC). *, p < 0.05. **, p < 0.01. ***, p < 0.001. ns, no significance. **d**. Hemolysis (HC_25_) on rat red blood cells and cytotoxicity (CC_50_) on HEK293T cells of ten peptides. **e**. Outer membrane permeabilization of ten peptides on *A.baumannii* ATCC19606 using *N*-phenyl-1-naphthylamine (NPN).

We tested minimal inhibitory concentrations (MICs), modes of action on membranes, hemolysis, and cytotoxicity of these ten peptides (Fig. 4a). We measured their MICs against two Gram-negative strains (*E. coli* ATCC25922 and *A. baumannii* ATCC19606) and two Grampositive strains (*S. aureus* ATCC6538 and *B. subtilis* ATCC6633). As shown in Fig. 4b, all ten peptides achieved at least an MIC ≤ 64 µM, demonstrating a success rate of 100% for being AMPs. This indicates that AMPainterV2 outperformed our previous AMPainter model^17^ in evolving AMPs from random sequences (success rate of 60% for AMPainter). Furthermore, eight peptides could inhibit all four bacterial strains, showing broad-spectrum antibacterial potency. Peptide V2-R08 exhibited a high level of antimicrobial activity with a mean MIC of 1.5 µM, while V2-R05 also had a mean MIC of 2.25 µM.

We utilized a fluorescent dye, propidium iodide (PI), to monitor the disruption of the inner membrane of bacteria under AMP treatment for Gram-positive *S. aureus* and Gram-negative *A. baumannii* (Fig. 4c). PI can interact with nuclear chromatin and generate fluorescent signals when the cytoplasmic membrane is disrupted, making it a commonly used method to detect the membrane-lytic mechanism of AMPs. We used a well-known membrane-lytic antibiotic, polymyxin B, as the positive control and the system without adding antimicrobial agents as the negative control (NC). Fig. 4c shows that all the evolved AMPs have membrane-lytic abilities against *S. aureus*. Notably, they produced stronger fluorescent signals than polymyxin B. On *A. baumannii*, nine evolved AMPs and polymyxin B exhibited membrane-lytic activity compared with NC, while the non-AMP V2-R10 showed no significant fluorescent signals. V2-R06, V2-R07, and V2-R08 were superior to polymyxin B in disrupting the inner membranes.

Considering that membrane-lytic AMPs are likely to disrupt human cell membranes as well, we evaluated their hemolysis on rat red blood cells and cytotoxicity on HEK293T cells. Fig. 4d displayed their HC_25_ and CC_50_ values. Seven out of ten peptides have a low hemolytic risk with HC_25_ > 128 µM, except for V2-R04, V2-R06, and V2-R10. We calculated the selectivity index (SI, CC_50_/mean MIC) in Table S4, where only V2-R04 had an SI of less than one. Most AMPs exhibited good selectivity towards bacterial membranes rather than human membranes, particularly V2-R05 (SI = 35.06) and V2-R08 (SI = 45.89).

We also examined their outer membrane permeabilization on Gram-negative *A. baumannii* via the *N*-phenyl-1-naphthylamine (NPN) uptake assay (Fig. 4e). All ten peptides effectively permeated the outer membrane of *A. baumannii*, with signals well above the NC. Remarkably, V2-R10 caused NPN fluorescence nearly ten times higher than polymyxin B, yet it failed to inhibit the growth of *A. baumannii* at this concentration. This result suggests that V2-R10 is particularly effective at disrupting the outer membrane while leaving the inner membrane intact (Fig. 4c). Overall, the successful validation of all ten evolved peptides as functional membrane-lytic AMPs demonstrates the design capability of AMPainterV2.

### Miniaturize AMPs with Deletion-Only AMPainterV2

In the production and application of AMPs, shorter sequences are usually preferred due to their lower chemical synthesis costs. ^24^ A conventional strategy is to truncate AMPs while preserving the core active region. ^25^ However, this trial-and-error approach is often case-dependent. Similarly, in synthetic biology, shorter proteins are favored to circumvent engineering challenges associated with expression and delivery. To date, only a few tools, such as Raygun ^26^ and SCISOR, ^27^ have been developed to shorten proteins to ideal lengths while maintaining their functions. However, these tools were not trained on short peptides, and Raygun is further restricted to proteins of at least 50 residues. A key feature of AMPainterV2 is its support for insertion, deletion, and mutation operations. This flexibility allows users to prioritize particular edit types when designing AMPs. Therefore, we provided a deletion-only version of AMPainterV2 that activates only the expert subnetwork responsible for deletion, specifically aiming to miniaturize AMPs.

We began by comparing the deletion-only AMPainterV2 with SCISOR, a discrete diffusion model. As AMPainterV2 was trained on membrane-lytic AMPs and SCISOR was trained on UniRef90, we used the recent AMP dataset again to avoid potential data leakage. Inspired by the insertion noising process during SCISOR’s training, we inserted random residues into these AMPs at random sites until the sequences were extended by 50%. These lengthened sequences were then fed into either the deletion-only AMPainterV2 or SCISOR for deletion (Fig. 5a). We anticipated that the models would delete the inserted noise residues to recover the original AMPs or yield peptides with improved antimicrobial activity. Fig. 5b displayed the self-consistent edit distance, similarity, and ΔHyperAMP score between each initial AMP and its corresponding deleted peptide. Compared to the three SCISOR variants (U90L, M, and S), AMPainterV2 exhibited lower edit distances and higher similarities. Nearly all peptides processed by SCISOR showed a decline in antimicrobial activity (ΔHyperAMP<0), whereas the average ΔHyperAMP score for the AMPainterV2 deleted peptides was −0.018, indicating that our deletion-only model can preserve activity. In fact, several peptides even exhibited increasing ΔHyperAMP scores greater than 0.5, suggesting that AMPainterV2 can identify superior candidates during this deletion process.

**Figure 5:**
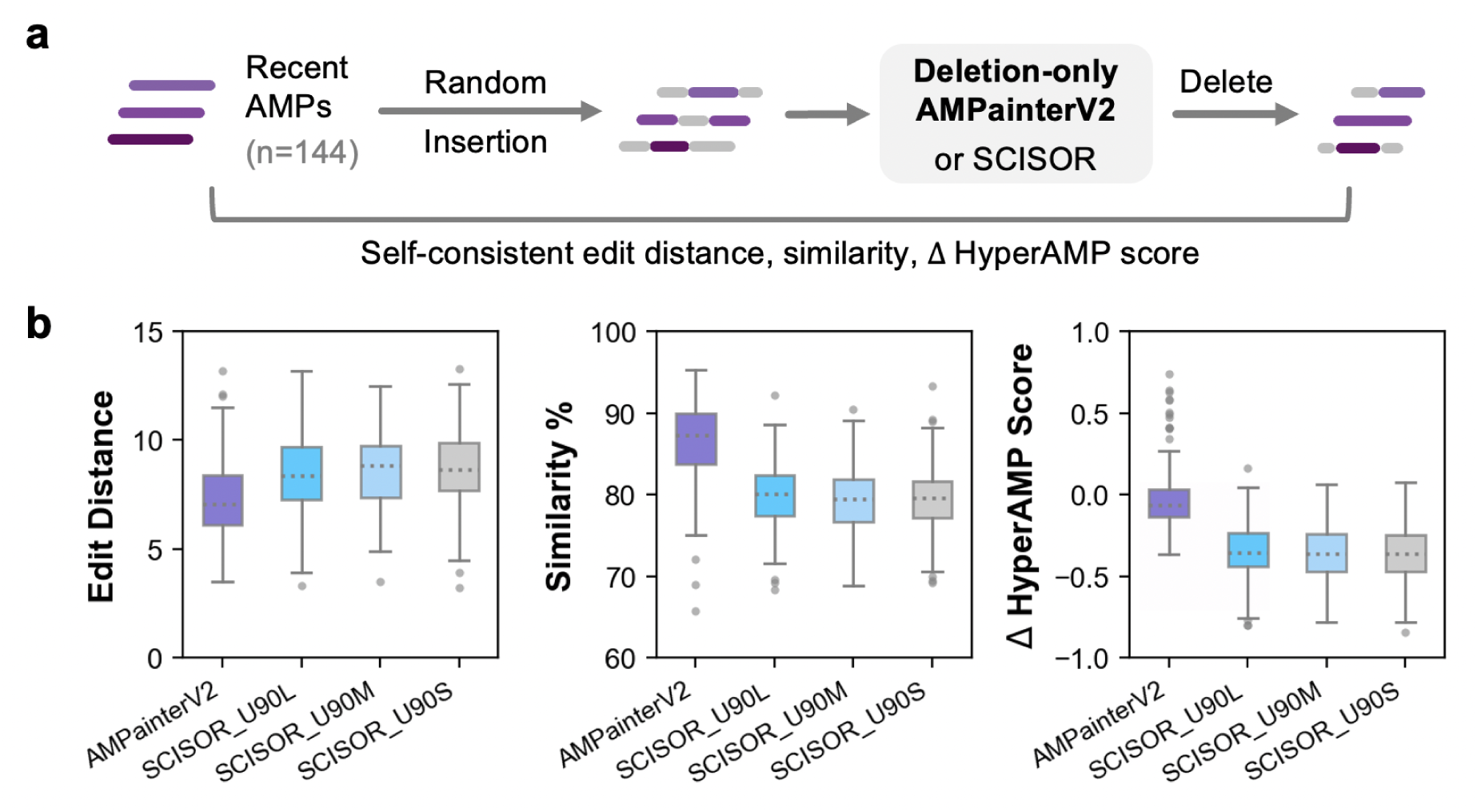
Comparison of the deletion-only AMPainterV2 and SCISOR. **a**. Comparison workflow using recent AMP sequences. **b**. Results on self-consistent edit distance, similarity, and the change in antimicrobial activity score.

We applied the deletion-only AMPainterV2 to the ten evolved membrane-lytic AMPs characterized in Fig. 4, along with the R04 peptide designed by AMPainter (hereafter called V1-R04). As these 11 peptides each exceed 20 amino acids in length, we aimed to miniaturize them by performing ten consecutive deletion rounds (Fig. 6a). All sequences converged after ten rounds, but the antimicrobial activity predicted by HyperAMP showed a noticeable decline as more residues were deleted (Table S5). As we aimed to maintain antimicrobial activity while achieving maximal length reduction, we selected the shortened peptide with the minimal value of |Δ*HyperAMPscore*|/#*Deletions* as the miniaturized peptide for each AMP. These miniaturized peptides were designated with a ‘-D’ suffix and synthesized for experimental validation.

**Figure 6:**
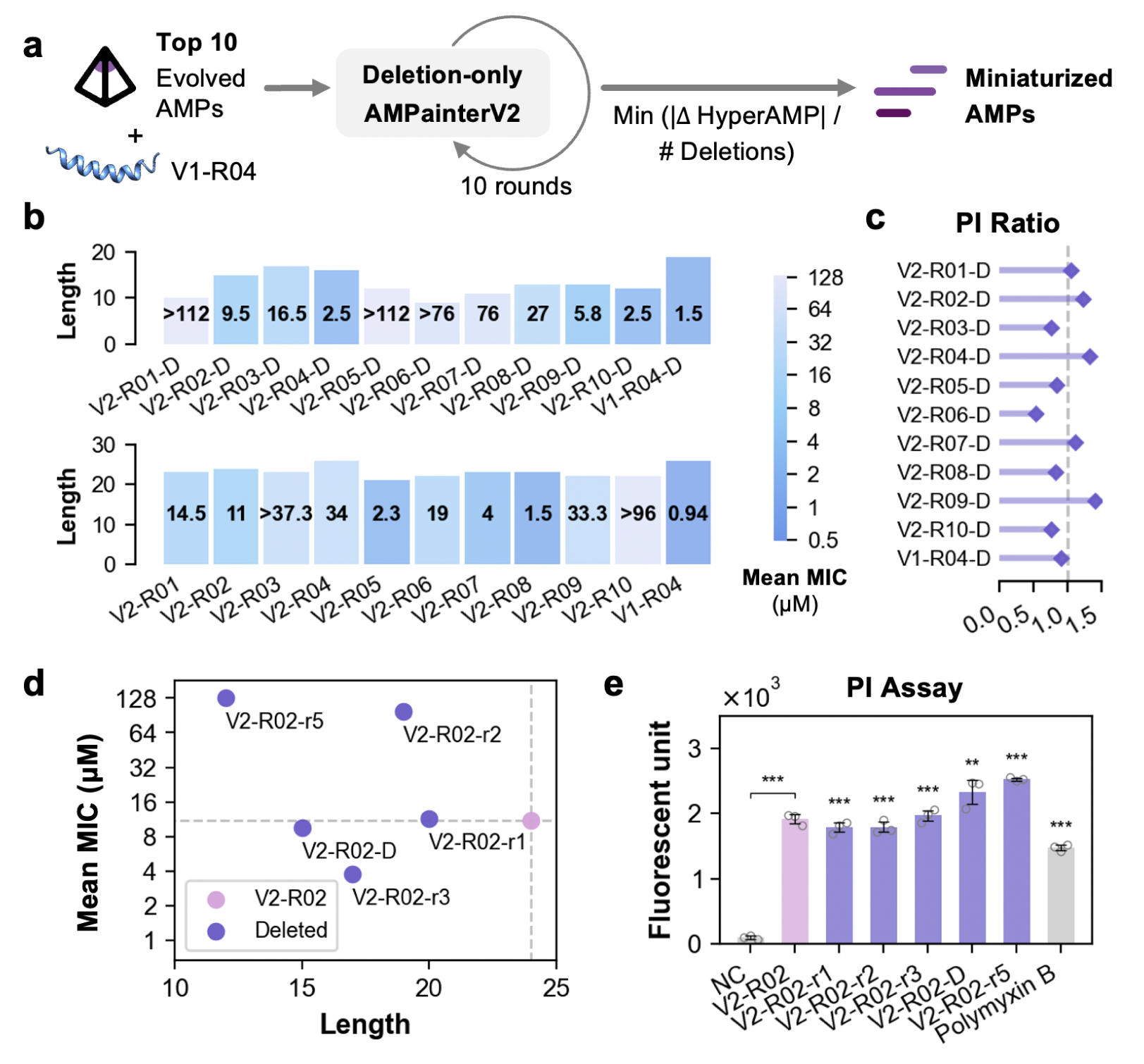
Miniaturization of evolved AMPs with deletion-only AMPainterV2. **a**. Workflow. **b**. Lengths and mean MICs of miniaturized AMPs in comparison to their original AMPs. **c**. Membrane-lytic ability (represented as PI ratio) of miniaturized AMPs in comparison to their original AMPs. **d**. Mean MICs versus sequence length of V2-R02 shortened peptides. **e**. Results of PI assay on *S.aureus* of V2-R02 shortened peptides. Unpaired t-tests were used for comparison with the negative control (NC). **, p < 0.01. ***, p < 0.001.

Fig. 6b shows that 11 miniaturized AMPs were shortened by approximately 10 residues, corresponding to an average length reduction of 42.7%. Six of them exhibited reduced or comparable mean MIC values against four bacterial strains, relative to their original AMPs. This strategy successfully miniaturized several highly active AMPs while maintaining their activity (e.g., V2-R02 and V1-R04), although it didn’t keep the activity level for V2-R05, V2-R07, and V2-R08. In some cases, miniaturization even led to marked activity enhancement. For instance, V2-R10 had a mean MIC of >96 µM while its miniaturized peptide V2-R10-D exhibited 2.5 µM. In the aspect of membrane-lytic mechanism, we also performed the PI assay on *S. aureus* for miniaturized peptides. As shown in Fig. 6c, five miniaturized peptides exhibited PI fluorescence ratios greater than one, indicating enhanced membrane disruption compared to their corresponding original AMPs. The remaining miniaturized peptides displayed PI ratios around 0.8 (except for V2-R06-D), suggesting that their membrane-lytic ability was largely preserved.

We examined our selection rule for miniaturized peptides using V2-R02 and V2-R04. Over ten deletion rounds, V2-R02 yielded five shortened intermediates, with 4, 5, 7, 9, and 12 residues deleted, respectively (Table S5). Three of them retained comparable or improved mean MIC values (Fig. 6d), and all five exhibited stronger membrane-lytic activity than polymyxin B (Fig. 6e). Among them, the selected miniaturized peptide, V2-R02-D, was the shortest peptide that pre-served both antimicrobial activity and membrane-lytic ability, consistent with our miniaturization objective. Interestingly, V2-R02-r5 showed the highest PI fluorescence (Fig. 6e). However, its MIC against *S. aureus* was >128 µM (red circle in Fig. S5a). Given that the PI assay was performed at a concentration of 512 µM, we speculated that V2-R02-r5 might inhibit *S. aureus* at higher concentrations. Further MIC measurements revealed that its MIC should be around 1024 µM (Fig. S5b). 128, 256, and 512 µM V2-R02-r5 also showed slight bacterial inhibition, which agrees with the observed membrane-lytic activity.

As for V2-R04, our model maintained its sequence pattern of alternating hydrophobic and hydrophilic residues, such as ‘KI’ and ‘RI’ motifs after deletions (Fig. S6a). However, in vitro MICs of these shortened peptides apparently increased when the sequence length was less than 10 residues (Fig. S6b). This trend could be captured by HyperAMP scores, which excluded them from being selected as the miniaturized peptide. The selected V2-R04-D exhibited the best activity among six shortened peptides. Notably, all six shortened peptides of V2-R04 demonstrated good membrane-lytic ability, surpassing both V2-R04 and polymyxin B (Fig. S6c). Overall, our miniaturization strategy successfully balanced functional preservation with length reduction. The deletion-only AMPainterV2 thus provides a powerful tool to miniaturize AMPs, facilitating their practical application.

## Discussion

In this study, we propose AMPainterV2, a model that designs membrane-lytic AMP sequences through insertion, deletion, and mutation operations. This work makes three main contributions. First, from a technical standpoint, AMPainterV2 addresses a gap in discrete optimization models by accommodating variable-length operations (insertion and deletion). We lever-aged an MoE architecture to explicitly assign distinct sequence edit operations to specialized expert subnetworks. Insertions and deletions played roles in enhancing the activity of AMPs (Fig. 2b) and enabled AMPainterV2 to outperform other discrete and continuous AMP optimization models (Fig. 3).

Second, from a practical perspective, AMPainterV2 designed membrane-lytic AMPs in a mechanism-driven manner. Membrane disruption confers broad-spectrum antimicrobial activity to AMP and reduces the likelihood of inducing resistance. However, previous studies on AMP generation either overlooked their mechanisms or relied on arbitrary physicochemical rules. By integrating the evolutionary design strategy with GAIL, our model successfully imitated the patterns of known membrane-lytic AMPs used as demonstration data. In experimental validation, all ten evolved peptides were confirmed to function as membrane-lytic AMPs, and some were highly active. Furthermore, the GAIL training framework is readily generalizable to other peptide classes with specialized mechanisms, simply by replacing the demonstration data accordingly.

Third, building on these two points, the deletion-only version of AMPainterV2 offers a useful tool for miniaturizing AMPs while preserving or even enhancing their antimicrobial potency and membrane-lytic mechanism, which have also been validated in vitro. This helps to save production costs and mitigate potential issues associated with longer sequences.

Despite its advances in designing membrane-lytic AMPs, AMPainterV2 has several limitations. First, the model is restricted to canonical peptide sequences with lengths ranging from 5 to 40 amino acids, aligning with AMPainter and HyperAMP. ^17^ AMPs containing noncanonical amino acids or chemical modifications still await exploration. Second, the number of demonstration membrane-lytic AMPs is limited, as we only included sequences with experimentally validated mechanisms. As we continue to update MemAMPdb, ^19^ more reported membrane-lytic AMPs will be incorporated. Finally, AMPainterV2 cannot yet evolve any random peptide into a membrane-lytic AMP within limited steps, which could potentially be addressed through iterative design or by expanding training sequences.

Recent advances in AMP mining and generation have significantly enriched the AMP repository. ^6,7,23^ Nevertheless, many novel AMPs lack mechanistic characterization, which hinders their practical application. As the success rate for discovering general AMPs has already been very high (e.g., 100% for 100 tested peptides ^28^), we believe the next frontier lies in designing AMPs with specialized functions, such as cell-penetrating AMPs for intracellular infections. ^29^ Our GAIL-based framework offers a powerful and generalizable approach that can be adapted to other peptides classes of interest, requiring small-scale data and minimal additional filtering.

## Materials and Methods

### AMPainterV2 model

#### MoE policy network

The mixture-of-experts (MoE) ^16^ architecture was adopted as the policy network to be compatible with the insertion, deletion, and mutation operations. The MoE receives the embedding of peptide sequences and produces the probability distribution over sequence edit operations (*p*(*Operation*)) and the residue sites (*p*(*Residue*)). As shown in Fig. 1, MoE policy network is composed of a router and three expert subnetworks. The router is a single fully connected layer with a softmax function, which transforms the embedding sequence *X* ∈ **R***^L^*^×^*^N^* to *p*(*Operation*) ∈ **R***^L^*^×3^ (Eq. 1):

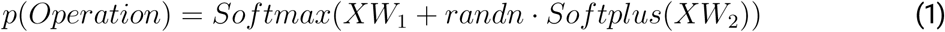

where *L* denotes the sequence length, *N* is the embedding dimension (128 was used here), *W*_1_ and *W*_2_ are trainable weight matrices, *randn* provides a random number following the standard Gaussian distribution, and *Softplus(x)* = *log*(1 + *e^x^*). Random noise was injected to balance the activation of different experts.

Based on *p*(*Operation*), the expert with the highest probability was selected for the up-coming evolution. Three experts were set to represent the three sequence operations: insertion, deletion, and mutation. Each expert subnetwork is a 2-layer neural network, which imports the sequence embedding *X* and exports the selection weight *E_i_*(*X*) of its representative operation *i*. The final probability of assigning amino acid sites is the weighted sum of each expert output:

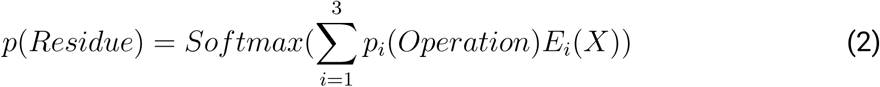

The assigned residue site is sampled according to *p*(*Residue*). Assuming that *A_L_*_−1_ was selected by MoE in an iteration for a peptide sequence *A*_0_*A*_1_*…A_L_*_−2_*A_L_*_−1_*A_L_*, the sequence would become *A*_0_*A*_1_*…A_L_*_−2_ *< mask > A_L_*_−1_*A_L_*for the insertion operation, *A*_0_*A*_1_*…A_L_*_−2_ *< mask > A_L_* for the mutation operation, or *A*_0_*A*_1_*…A_L_*_−2_*A_L_*for the deletion operation. The string with the *< mask >* token would be fed into the fine-tuned language model in the next stage to decode the masking residue. That is to say, insertion means inserting a new residue before the assigned residue, mutation means replacing the assigned residue with a new residue, and deletion means removing the assigned residue. A constraint on the length range was set for these variable-length operations. If the sequence length is less than 5 or greater than 40 amino acids after editing, this operation will not be accepted, and the sequence will remain unchanged.

#### Discriminator

The discriminator took the peptide sequence as input and produced the probability of it being a membrane-lytic AMP. Four hundred eighty-one membrane-lytic AMPs from MemAMPdb, ^19^ whose lengths are less than 40, were taken as positive samples, and the peptides designed by the AMPainterV2 agent were labeled as negative. In each episode, all designed peptides were placed in the buffer, and positive sequences were randomly selected to fill the buffer to its maximum capacity of 256 sequences.

The sequences were embedded with 531 physicochemical properties from AAindex. ^30^ The peptide embeddings were projected to the query *Q*, key *K*, and value *V* with linear layers, and then processed with the agent attention module (Eq. (3)). ^31,32^ Agent attention introduced agent tokens *A* in addition to conventional query *Q*, key *K*, and value *V* in attention mechanisms, serving as a bridge to integrate softmax and linear attention, making it more effective and cost-efficient. ^31^ Following the default setting, *A* = *pooling*(*Q*) was used.

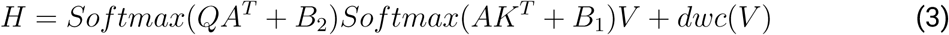

where *B*_1_ and *B*_2_ are the agent bias, and *dwc* refers to a depthwise convolution module.

After obtaining the 128-dimensional vector *H* from the agent attention module, a two-layer neural network was used to produce the probabilities. The loss function of the discriminator was binary cross entropy. Since the discriminator weights were randomly initialized, a linear learning rate warm-up strategy was adopted for 10 steps. The highest learning rate was 0.0005. The batch size was set to 64 and the optimizer was Adam with a weight decay of 0.0005. The training epoch was 10 in each episode.

#### GAIL training

The MoE policy network was trained using generative adversarial imitation learning (GAIL) ^18^ with membrane-lytic AMPs. Like AMPainter, ^17^ the policy network was used to assign the site (and edit operations here), and a fine-tuned language model was used for decoding the new residue. The language model Ankh fine-tuned with AMPs in AMPainter was also used in this work. ^17^ GAIL was used to train the MoE policy network. The training objective of GAIL can be defined as Eq. (4), to find a balance between the policy *π* and the discriminator *D*:

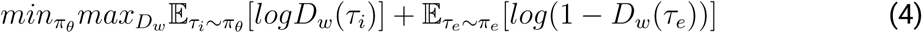

where *π_θ_* is the policy with trainable parameter *θ*, *D_w_* is the discriminator with trainable parameter *w*. *π_e_* is the expert policy and *τ_e_* is the expert trajectory, *τ_i_* is the trajectory from policy *π_θ_*.

For the discriminator *D*, the weights *w* were updated using the gradient descent-based Adam optimizer. The policy *π* was trained using the policy gradient-based REINFORCE algorithm. ^33^ Define the trajectory *τ* = {*s*_1_*, a*_1_*, r*_1_*, …, s_t_, a_t_, r_t_*} at timestep *t*, the state *s_t_* is the peptide sequence, action *a_t_* is the assigned residue and correlated operation as defined by *π_θ_*, and the reward *r_t_* is the output probability of *D_w_*. During *T* − 1 iterations of an episode, *G_t_* is the sum of rewards from each iteration, and *J* (*θ*) is the objective function to update *θ*:

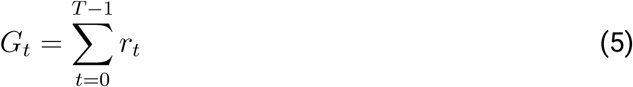

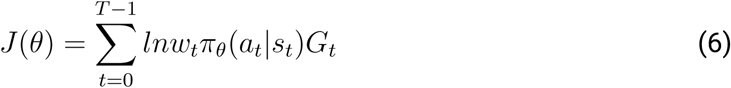

where *w_t_* is a discounting weight of *π_θ_*(*a_t_*|*s_t_*), which was set as the end condition or punishment for repeated sequences(Eq. (7)).

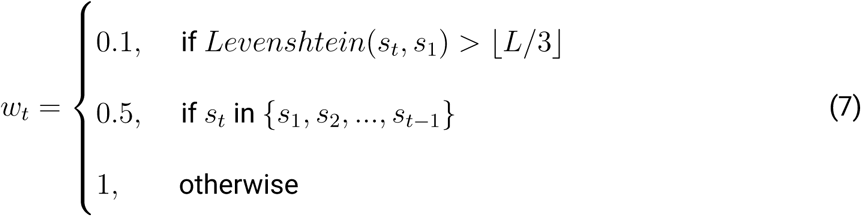

966 random sequences with lengths ranging from 10 to 40 were used as the training dataset. ^17^ To build it, 1000 peptide sequences were randomly generated first, and the cd-hit ^34^ tool was used to remove redundancy with an identity cutoff of 0.7. The maximum number of training episodes was set to 40. The AMPainterV2 model was trained on an NVIDIA A40 GPU. The optimizer was Adam, and the learning rate was 0.001. The batch size was set to 64.

Two additional evaluation metrics were used to monitor the training process, perplexity and Boman index. In each episode, the average score of designed peptides was calculated (Fig. S1). Perplexity is commonly used to evaluate the quality of generated protein sequences. A lower perplexity score indicates a more protein-like sequence, as recognized by large language models. Here, this score was implemented with the ESM-2 8M model. ^35,36^

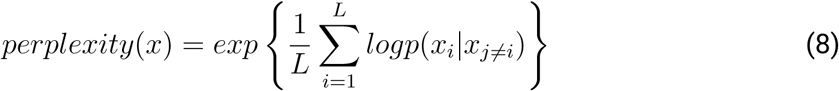

where *L* is the length of the input sequence.

Boman index was used to estimate the membrane-interacting propensity of peptides. ^37^ A low value of Boman index may indicate that the peptide can interact with membranes. Boman index was calculated using the modlAMP package. ^38^

#### Inference

The weights of policy network with the highest HyperAMP score during training (episode 30) were adopted. At the inference stage, only the agent of AMPainterV2 was used, including the MoE policy network and the fine-tuned language model. AMPainterV2 took in a batch of input sequences simultaneously. The episode was set to 10 to evolve 10 parallel paths, and the iteration was assigned a maximum of 13. All sequences were collected and ranked using the HyperAMP model. ^17^

### Comparison with other models

Four published models for AMP optimization were used as baselines here, including EvoGradient, ^15^ PepZOO, ^12^ deepAMP, ^20^ and HydrAMP. ^11^ Among them, AMPainter and EvoGradient are fixed-length discrete optimization methods, while the other three are continuous optimization methods.

1. ApexGO: ^10^ a VAE model using Bayesian optimization with Gaussion process in the la-tent space for AMP optimization, guided by predicted minimal inhibitory concentration scores against multiple bacteria from APEX predictor.
2. EvoGradient: ^15^ a gradient-based virtual directed evolution method for AMPs. During the back-propagation process of their AMP-READ predictor, a gradient descent with projection was adopted to optimize the input sequence with discrete perturbations.
3. PepZOO: ^12^ an AMP optimization method utilizing a zeroth-order optimization approach in the continuous latent space compressed by a VAE model. The model for optimizing antimicrobial function and activity was used.
4. deepAMP: ^20^ a VAE model to generate AMP analogs with enhanced activity. Three training steps were incorporated, including pretraining on UniProt peptides, fine-tuning on known AMP degeneration pairs, and on the analog pairs of target AMP. The model that completed the second training stage (deepAMP-AOM) was used here for comparison, because we require to evolve hundreds of sequences with one model, and we cannot fine-tune the model for each sequence.
5. HydrAMP: ^11^ a conditional VAE model for AMP generation. Here, its analog generation task with discovery mode was adopted, and the default filters were included.

Two initial datasets were used for comparison, including an AMP dataset and a random sequence dataset. For the AMP dataset, 199 AMP sequences were collected from articles published in the past two years. These AMPs met two criteria: (i) They have been experimentally validated with at least one MIC less than 128 µM or µg/mL; (ii) Their antimicrobial mechanisms have not been reported. 144 AMPs, whose lengths were less than 25 amino acids, were retained to meet the length requirement of HydrAMP and PepZOO. For random sequences, the first 1000 sequences with lengths ranging from 5 to 25 residues were randomly generated. Then, cd-hit was used to remove similar sequences with a threshold of 0.7. 200 sequences were randomly selected as the dataset. The maximum length of 25 was set due to the restrictions of PepZOO and HydrAMP. All sequences were ensured not to overlap with sequences in the training set of each method.

The server of HydrAMP was used, and other methods were implemented with their source codes locally. All the parameters and settings were their defaults. Ten parallel iterations were run for each input sequence. Only evolved sequences whose Levenshtein distance from its initial peptide was less than a third of the initial length were kept. As for ApexGO optimization, the same similarity constraint was applied to each peptide as template. The number of random initialization sequences was 2000, the maximum number of oracle calls was limited to 2000, and the number of output optimized sequences was fixed at 10. The optimization objective was defined as the average score across all bacterial strains.

The top 100 (for the AMP dataset) or 200 (for the random dataset) of these evolved sequences from each method, evaluated by HyperAMP score, were collected for further evaluation and comparison. In the analysis of non-toxic AMPs, BioToxinPept ^6^ and ToxinPred 3.0^21^ were utilized for toxicity assessment. The standards for non-toxic AMP were a HyperAMP score greater than 1.0 (for the AMP dataset) or 0.5 (for the random dataset), along with a BioToxinPept probability of less than 0.5, or a ToxinPred 3.0 probability of less than 0.38.

The diversity score was calculated using pairwise Levenshtein distance within the sequence set, similar to AMPainter. ^17^

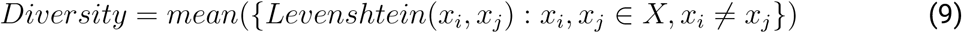

where *X* is the output set of sequences, and *x_i_*and *x_j_* are two distinct sequences.

Local alignment based on the Smith-Waterman algorithm was adopted to calculate the pairwise similarity of each evolved sequence and membrane-lytic AMP using the BLOSUM62 matrix. An average score was reported for each sequence set.

### Model variant trained with non-specific AMPs

To construct a non-specific AMP set without assigned mechanisms, 19,670 AMP sequences from the training set of AMP-Diffusion ^39^ were used. Sequences with lengths ranging from 10 to 40 residues were retained and de-duplicated using cd-hit ^34^ with a cut-off of 0.7. The inter-dataset redundancy with membrane-lytic AMPs was then removed using cd-hit-2d with a stricter cut-off of 0.5, resulting in 3,331 sequences. To ensure a fair comparison, 481 sequences were extracted that had similar length and amino acid distributions to the membrane-lytic AMPs. The model settings and training procedure are the same as those of AMPsainterV2, with the only change being the replacement of membrane-lytic AMPs with these non-specific AMPs as demonstration data.

### Deletion-only AMPainterV2

AMPainterV2 with user assigned operations was provided at the inference stage. As for the deletion-only version, only the deletion operation was allowed after selecting the expert according to *p(Operation).* If insertion or mutation was selected, the sequence would not change at this step. Additionally, only the deletion expert network could be activated when calculating *p(Residue)*.

To compare the deletion-only AMPainterV2 with SCISOR, the AMP dataset containing 144 recently reported AMPs with lengths of less than 25 amino acids was used as the initial data. Random amino acids were inserted into the AMP sequences. For each AMP, ten different randomly inserted sequences were produced. The number of inserted amino acids was half the length of each initial AMP sequence. Therefore, the maximum length of inserted sequences was 38, satisfying the length limit (40 residues) of AMPainterV2. 1,440 inserted sequences were deleted by deletion-only AMPainterV2 for one episode, as well as three versions of SCISOR (small, medium, and large models, trained on UniRef90 data) with a shrinking per-centage of 30%. The Levenshtein edit distance, similarity, Δ HyperAMP score were calculated between each pair of initial AMP and its shortened sequence. The average score was reported on ten samples for each initial AMP.

To miniaturize the evolved membrane-lytic AMPs, 11 peptides (V2-R01∼V2-R10 and V1-R04) were evolved for ten rounds using deletion-only AMPainterV2. In each round, ten parallel evolutions and 13 iterations of operations were performed. The sequence with the highest HyperAMP score was kept among the ten final sequences from parallel evolutions and entered the next round. The number of deletions (# Deletions) and the change in HyperAMP scores between each shortened sequence and its original sequence (Δ HyperAMP score) were calculated. The shortened sequence with minimal |Δ HyperAMP score|/# Deletions was selected as the miniaturized peptide for experimental validation.

### Wet-lab experiments

#### Peptide synthesis

All peptides were synthesized via solid-phase peptide synthesis by DGpeptide Co., Ltd. Their molecular weights were verified by mass spectrometry, and their purity (>95%) was confirmed by HPLC (Fig. S7 - Fig. S41).

#### MIC measurement

Four standard bacteria used for the MIC test were *Escherichia coli* ATCC25922, *Acinetobacter baumannii* ATCC19606, *Staphylococcus aureus* ATCC6538, and *Bacillus subtilis* ATCC6633. The testing procedures followed the Hancock methods. ^40^ The peptides were diluted in sterilized water containing 5% dimethyl-sulfoxide (DMSO) to 512 µM. Then a two-fold gradient dilution was performed to create different concentrations (128, 64, 32, 16, 8, 4, 2, 1, 0.5 µM) of AMPs. Each bacteria was incubated in Luria-Bertani (LB) broth at 37°C until its absorbance at 625 nm reached 0.1, then diluted 10,000 times and added into a sterile 96-well polypropylene plate (Grenier). A co-incubation system included 100 µL peptide solution and 100 µL bacterial suspension. After incubation at 37°C for 18 hours, the MIC was determined as the lowest pep-tide concentration that prevented bacterial growth. All tests were performed in triplicate wells, and the MIC readings from the triplicate wells were consistent.

#### Hemolysis assay

*In vitro* hemolysis and cytotoxicity were measured by WuXi AppTec Co., Ltd. The rat blood was collected and mixed in PBS, then centrifuged at 500 g for 5 min to make the red blood cell solution (RBC, 10%). Two-fold gradient dilution was performed to obtain different concentrations (128, 64, 32, 16, 8, 4, 2, and 1 µM) of AMP solutions. Then, AMP solutions were added to 100 µL RBC to make the test solution and incubated at 37°C for 1 hour. The assay solution was centrifuged at 2500 g for 6 min. After centrifugation, the 75 µL supernatant was placed into a 96-well plate. PBS was used as the vehicle control, and 0.1% Triton X-100 served as the positive control. The absorbance of the supernatant in each well was measured at 450 nm using EnVision (PerkinElmer), and the percentage of hemolysis was calculated as Eq. (10). The HC_25_value was fitted using GraphPad Prism 8. All tests were performed in triplicate.

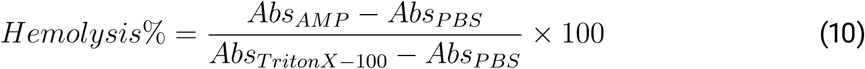

#### Cytotoxicity assay

HEK293T cells were cultured in DMEM supplemented with 10% FBS and incubated at 37°C with 5% CO_2_. After planting the cells into a 96-well plate and incubating overnight, AMP solutions were added. The gradient dilution of AMPs followed the same settings as mentioned in the hemolysis assay. Staurosporine (STS) was used as a positive control and PBS as a vehicle control. STS was diluted to 60, 12, 2.4, 0.48, 0.1, 0.02, 0.0038, and 0.0008 µM. After incubation for 72 hrs, the CellTiter-Glo® Luminescent assay was performed to evaluate cell viability. After equilibration to room temperature, the 96-well plate containing treated cells was loaded with 100 µL CellTiter Glo (CTG, Promega) reagent in each well. The plate was incubated for 30 min, then detected using the EnVision system at a wavelength of 570 nm. Data were fitted and analyzed with GraphPad Prism 8. All tests were performed in triplicate.

#### Inner membrane disruption assay

*S. aureus* ATCC6533 or *A. baumannii* ATCC19606 cells were cultured in LB broth to the exponential stage and then centrifuged at 4000 g. After discarding the supernatant, the precipitated bacterial cells were washed with PBS three times and resuspended in PBS to obtain OD_600_ = 1.0. Using a black 96-well polypropylene plate, 10 µL bacterial suspension was mixed with 10 µL AMP solutions in PBS to reach a final concentration of 4×MIC. For peptides whose MIC values are larger than 128 µM, a final concentration of 512 µM was used. After incubation for 1 h at 37°C, 80 µL propidium iodide (PI, Solarbio, #C0080) solution was added at a final concentration of 20 µM and incubated for another 40 min in the dark. Fluorescent signals were measured under the excitation of 535 nm and emission of 615 nm using a BioTek Cytation5 plate reader. The positive control was polymyxin B, while the negative control (NC) was composed of PBS, bacteria, and PI. The final concentration of polymyxin B was also 4×MIC. The fluorescent values were corrected by subtracting the blank control (20 µL PBS and 80 µL PI). The PI ratio for comparing the miniaturized AMP and its original AMP was calculated as the quotient of their PI fluorescence. All samples were tested in triplicate.

#### Outer membrane permeabilization assay

*A. baumannii* ATCC19606 cells were cultured in LB broth to the exponential stage. The bacterial solution was centrifuged at 4000 g for 10 min to harvest, then washed with the buffer (5 mM HEPES containing 5 mM glucose, pH 7.2) three times and adjusted to OD_600_ = 0.5. *N*-phenyl-1-naphthylamine (NPN, Aladdin, P110559) was dissolved in acetone and added to the bacterial solution to reach a final concentration of 10 µM. The mixture solution of bacteria and NPN was incubated for 15 min at 37°C in the dark. 25 µL AMP solutions (conc. 4×MIC) were added to a black 96-well polypropylene plate, then 75 µL mixture solution was added to each well to make the final concentration of AMP 1×MIC. For peptides whose MIC values are larger than 128 µM, a final concentration of 128 µM was used. The positive control was polymyxin B, while the negative control (NC) was composed of buffer, bacteria, and NPN. Fluorescent signal was monitored every 5 min for 30 min using the BioTek Cytation5 plate reader. The excitation wavelength was 350 nm and the emission wavelength was 420 nm. The fluorescent values were corrected by subtraction of blank control (buffer and NPN at the same concentration). All samples were tested in triplicate.

## Data and code availability

All datasets and codes will be available after peer review at https://github.com/ComputBiophys/AMPainterV2.

## Author contributions

C.S. and R.D. conceived the project. R.D. conducted the research under the supervision of C.S.. R.D. wrote the original manuscript, and C.S. reviewed and edited the manuscript.

## Acknowledgements

The authors thank Zefeng Zhu and Yiechang Lin for their insightful suggestions and Qiushi Cao for assistance with the experiments. This work was supported by the National Key R&D Program of China (2024YFA0916800), the Innovative Research Group Project of the National Natural Science Foundation of China (T2321001), and the Frontier Innovation Fund of Peking University Chengdu Academy for Advanced Interdisciplinary Biotechnologies. The authors also thank the Core Facilities of Life Sciences at Peking University for providing the facilities.

