## Supplementary Information for "Evolutionary Design of Membrane-Lytic Antimicrobial Peptides with Mixture of Experts"

### Supplementary Tables

**Table S1:** Ratio of toxic AMPs in the training set for each method, as evaluated by BioToxinPept or ToxinPred 3.0.

| Model | AMPainterV2 | EvoGradient | PepZOO | deepAMP | HydrAMP |
|---|---|---|---|---|---|
| BioToxinPept | 14.97% | 21.88% | 27.98% | 24.04% | 31.39% |
| ToxinPred 3.0 | 35.14% | 45.52% | 38.37% | 42.51% | 48.05% |

**Table S2:** Comparison of evolving AMPs or random sequences using the AMPainterV2 model trained with membrane-lytic AMPs or non-specific AMPs.

| Dataset | Model (demonstration data) | Non-toxic AMP Ratio% | | SVM Pass% | Perplexity |
|---|---|---|---|---|---|
| | | ToxinPred 3.0 | BioToxinPept | | |
| AMP | AMPainterV2 (membrane-lytic AMPs) | 58.0 | 96.0 | 96.0 | 12.991 ± 3.724 |
| | AMPainterV2 (non-specific AMPs) | 60.0 | 96.0 | 81.0 | 12.171 ± 4.687 |
| Random | AMPainterV2 (membrane-lytic AMPs) | 67.0 | 99.0 | 53.5 | 19.845 ± 2.677 |
| | AMPainterV2 (non-specific AMPs) | 62.5 | 94.0 | 39.5 | 19.009 ± 2.860 |

**Table S3:** Sequences, HyperAMP scores, and physicochemical properties of the top 10 peptides evolved by AMPainterV2. MW, molecular weight. H, hydrophobicity (on the Eisenberg scale). μH, hydrophobic moment (on the Eisenberg scale).

| Name | Sequence | Length | HyperAMP | MW | Charge | H | μH | Boman |
|---|---|---|---|---|---|---|---|---|
| V2-R01 | RIKPIRKGFLFAGKTFWVIYKGR | 23 | 1.13236 | 2782.38 | 6.99 | -0.013 | 0.228 | 1.307 |
| V2-R02 | YRFKGIAIKTLWGGLGKVKLRLKR | 24 | 1.09449 | 2802.46 | 7.99 | -0.081 | 0.328 | 1.283 |
| V2-R03 | VFHAWMKFMKKLFKFTFKIKWSK | 23 | 1.09320 | 3007.76 | 7.03 | 0.087 | 0.363 | 0.414 |
| V2-R04 | KSLYTWVKKIKHRIYKMWIWFGKHIF | 26 | 1.06279 | 3438.20 | 7.07 | 0.067 | 0.470 | 0.727 |
| V2-R05 | TPKTWILKLAKLHQKVKLLIR | 21 | 1.05687 | 2528.18 | 6.03 | -0.019 | 0.397 | 0.733 |
| V2-R06 | WRGWWRIISRGLVYSVVSRYAR | 22 | 1.05063 | 2767.20 | 4.99 | -0.060 | 0.129 | 2.160 |
| V2-R07 | LKIHAKLHNFYFLIKGLTLKKKG | 23 | 1.05059 | 2711.35 | 6.07 | 0.083 | 0.059 | 0.343 |
| V2-R08 | IPGTKWALKLAKKHMKLMKMLIR | 23 | 1.04807 | 2736.55 | 7.03 | -0.008 | 0.515 | 0.521 |
| V2-R09 | IWNTIKRLPIWPWGTLRWPRLF | 22 | 1.03829 | 2850.42 | 3.99 | 0.132 | 0.349 | 0.879 |
| V2-R10 | GLMLRWHWFWKVTHHLLKWFIM | 22 | 1.03532 | 2966.62 | 3.11 | 0.343 | 0.256 | -0.318 |

**Table S4:** Hemolysis, cytotoxicity, and selectivity index (SI, $CC_{50}$/Mean MIC) of the designed AMPs. Mean MIC with '>' indicates that this AMP has at least a MIC value of >128 μM, excluding their SI from the calculation and showing as '/'.

| Name | Mean MIC (μM) | $HC_{25}$ (μM) | $CC_{50}$ (μM) | SI |
|---|---|---|---|---|
| V2-R01 | 14.5 | >128 | 16.47 | 1.14 |
| V2-R02 | 11 | >128 | 127.5 | 11.59 |
| V2-R03 | >37.33 | >128 | 68.58 | / |
| V2-R04 | 34 | 10 | 22.9 | 0.67 |
| V2-R05 | 2.25 | >128 | 78.88 | 35.06 |
| V2-R06 | 19 | 40 | 78.39 | 4.13 |
| V2-R07 | 4 | >128 | 30.56 | 7.64 |
| V2-R08 | 1.5 | >128 | 68.83 | 45.89 |
| V2-R09 | 33.25 | >128 | 69.69 | 2.10 |
| V2-R10 | >96 | 30 | 27.46 | / |

**Table S5:** Miniaturized sequences and scores during ten rounds of deletion by the deletion-only AMPainterV2.

| Round | Name | Sequence | Length | HyperAMP | #Del | \|ΔHyperAMP\| /#Del × 100 | MIC (μM) | | | | E-value[a] | E-value[b] |
|---|---|---|---|---|---|---|---|---|---|---|---|---|
| | | | | | | | Ec | Ab | Sa | Bs | | |
| 1 | V2-R01-r1 | RIKIRKLFAGKFVIKR | 16 | 1.0325 | 7 | 1.4257 | 128 | 64 | 32 | 128 | / | / |
| 2 | V2-R01-r2 | RIKIKLFAVKR | 11 | 0.8359 | 12 | 2.4700 | | | | | / | / |
| 3 | **V2-R01-D** | RIKIKLAVKR | 10 | 0.9668 | 13 | **1.2738** | >128 | >128 | 64 | >128 | / | / |
| 4 | V2-R01-r4 | RIKIKLAKR | 9 | 0.8438 | 14 | 2.0614 | | | | | / | / |
| 5 | V2-R01-r4 | RIKIKLAKR | 9 | 0.8438 | 14 | 2.0614 | | | | | / | / |
| 6 | V2-R01-r4 | RIKIKLAKR | 9 | 0.8438 | 14 | 2.0614 | | | | | / | / |
| 7 | V2-R01-r7 | RIKIKLAR | 8 | 0.7938 | 15 | 2.2567 | | | | | / | / |
| 8 | V2-R01-r7 | RIKIKLAR | 8 | 0.7938 | 15 | 2.2567 | | | | | / | / |
| 9 | V2-R01-r9 | RIKILAR | 7 | 0.7945 | 16 | 2.1119 | | | | | / | / |
| 10 | V2-R01-r9 | RIKILAR | 7 | 0.7945 | 16 | 2.1119 | | | | | / | / |
| 1 | V2-R02-r1 | RKIAIKTLWGGLKVKLRLKR | 20 | 1.1894 | 4 | 2.3726 | 32 | 4 | 2 | 8 | / | 9.0 |
| 2 | V2-R02-r2 | RKIAIKLWGGLKVKLRLKR | 19 | 1.1876 | 5 | 1.8621 | >128 | >128 | 128 | >128 | / | 8.1 |
| 3 | V2-R02-r3 | RKAIKLWGLKVKLRLKR | 17 | 1.1265 | 7 | 0.4570 | 8 | 2 | 1 | 4 | / | 6.4 |
| 4 | **V2-R02-D** | RKAIKLWLKVLRLKR | 15 | 1.1039 | 9 | **0.1044** | 16 | 4 | 2 | 16 | / | 1.6 |
| 5 | V2-R02-r5 | RAIKLWLLRLKR | 12 | 1.0545 | 12 | 0.3333 | 128 | >128 | >128 | >128 | / | 3.0 |
| 6 | V2-R02-r5 | RAIKLWLLRLKR | 12 | 1.0545 | 12 | 0.3333 | 128 | >128 | >128 | >128 | / | 3.0 |
| 7 | V2-R02-r5 | RAIKLWLLRLKR | 12 | 1.0545 | 12 | 0.3333 | 128 | >128 | >128 | >128 | / | 3.0 |
| 8 | V2-R02-r5 | RAIKLWLLRLKR | 12 | 1.0545 | 12 | 0.3333 | 128 | >128 | >128 | >128 | / | 3.0 |
| 9 | V2-R02-r5 | RAIKLWLLRLKR | 12 | 1.0545 | 12 | 0.3333 | 128 | >128 | >128 | >128 | / | 3.0 |
| 10 | V2-R02-r5 | RAIKLWLLRLKR | 12 | 1.0545 | 12 | 0.3333 | 128 | >128 | >128 | >128 | / | 3.0 |
| 1 | **V2-R03-D** | HAWMKKKLKFTFKIKWK | 17 | 1.0907 | 6 | **0.0412** | 16 | 16 | 2 | 32 | / | / |
| 2 | V2-R03-r2 | HAWMKKLFTKIKWK | 14 | 0.9334 | 9 | 1.7760 | | | | | / | / |
| 3 | V2-R03-r3 | HAWMKKLTKIKWK | 13 | 0.8334 | 10 | 2.5981 | | | | | / | / |
| 4 | V2-R03-r4 | HAKLKIKWK | 9 | 0.9061 | 14 | 1.3364 | | | | | / | 3.8 |
| 5 | V2-R03-r5 | AKLKIKWK | 8 | 0.8864 | 15 | 1.3787 | | | | | / | 3.1 |
| 6 | V2-R03-r6 | AKLKIKW | 7 | 0.8095 | 16 | 1.7725 | | | | | / | 2.4 |
| 7 | V2-R03-r7 | AKLIKW | 6 | 0.6885 | 17 | 2.3806 | | | | | / | 0.5 |
| 8 | V2-R03-r8 | AKLIW | 5 | 0.6584 | 18 | 2.4156 | | | | | / | / |
| 9 | V2-R03-r8 | AKLIW | 5 | 0.6584 | 18 | 2.4156 | | | | | / | / |
| 10 | V2-R03-r8 | AKLIW | 5 | 0.6584 | 18 | 2.4156 | | | | | / | / |
| 1 | V2-R04-r1 | KLTWKKIHRIKMWIWFKHIF | 20 | 1.1141 | 6 | 0.8556 | 64 | 8 | 4 | 32 | / | / |
| 2 | **V2-R04-D** | KLTWKKIRIKWIKHIF | 16 | 1.0616 | 10 | **0.0120** | 4 | 1 | 1 | 4 | / | / |
| 3 | V2-R04-r3 | KLTWKIRIKIKI | 12 | 1.0060 | 14 | 0.4053 | 8 | 4 | 2 | 8 | / | / |
| 4 | V2-R04-r3 | KLTWKIRIKIKI | 12 | 1.0060 | 14 | 0.4050 | 8 | 4 | 2 | 8 | / | / |
| 5 | V2-R04-r4 | KLKIRIKII | 9 | 1.0336 | 17 | 0.1718 | 64 | >128 | 64 | 64 | / | / |
| 6 | V2-R04-r6 | KLKIRIKI | 8 | 0.9507 | 18 | 0.6228 | 128 | >128 | 64 | 128 | / | / |
| 7 | V2-R04-r6 | KLKIRIKI | 8 | 0.9507 | 18 | 0.6228 | 128 | >128 | 64 | 128 | / | / |
| 8 | V2-R04-r8 | KLKIRII | 7 | 0.7329 | 19 | 1.7358 | 128 | 64 | 8 | 32 | / | / |
| 9 | V2-R04-r8 | KLKIRII | 7 | 0.7329 | 19 | 1.7358 | 128 | 64 | 8 | 32 | / | / |
| 10 | V2-R04-r8 | KLKIRII | 7 | 0.7329 | 19 | 1.7358 | 128 | 64 | 8 | 32 | / | / |
| 1 | V2-R05-r1 | TPKTWILKLAKLHKKLLIR | 19 | 1.1046 | 2 | 2.3869 | | | | | / | / |
| 2 | V2-R05-r1 | TPKTWILKLAKLHKKLLIR | 19 | 1.1046 | 2 | 2.3869 | | | | | / | / |
| 3 | V2-R05-r3 | TPKTWIKLALKKLLIR | 16 | 1.0428 | 5 | 0.2816 | | | | | / | / |
| 4 | V2-R05-r3 | TPKTWIKLALKKLLIR | 16 | 1.0428 | 5 | 0.2820 | | | | | / | / |
| 5 | V2-R05-r5 | TPKWIKLALKLLIR | 14 | 1.0234 | 7 | 0.4786 | | | | | / | / |
| 6 | V2-R05-r6 | TPKWIKLAKLLIR | 13 | 1.0492 | 8 | 0.0963 | | | | | / | 8.6 |
| 7 | **V2-R05-D** | TPKWIKLKLLIR | 12 | 1.0552 | 9 | **0.0189** | >128 | >128 | 64 | 128 | / | / |
| 8 | V2-R05-r8 | PKWIKLKLLIR | 11 | 1.0343 | 10 | 0.2260 | | | | | / | / |
| 9 | V2-R05-r8 | PKWIKLKLLIR | 11 | 1.0343 | 10 | 0.2260 | | | | | / | / |
| 10 | V2-R05-r10 | PKIKLKLLIR | 10 | 0.9985 | 11 | 0.5309 | 128 | >128 | 32 | 64 | / | / |
| 11 | V2-R05-r10 | PKIKLKLLIR | 10 | 0.9985 | 11 | 0.5309 | 128 | >128 | 32 | 64 | / | / |
| 1 | V2-R06-r1 | WRWWRIIRGLVVVRYAR | 17 | 1.1358 | 5 | 1.7034 | | | | | / | / |
| 2 | V2-R06-r2 | WRWWRIIRGLVVRAR | 15 | 1.1263 | 7 | 1.0805 | | | | | / | / |
| 3 | V2-R06-r3 | RWRIILVRAR | 10 | 1.0739 | 12 | 0.1941 | | | | | / | 8.2 |
| 4 | **V2-R06-D** | RWRIIVRAR | 9 | 1.0478 | 13 | **0.0215** | 128 | >128 | 16 | 32 | / | / |
| 5 | V2-R06-r5 | RWRIIRAR | 8 | 0.8702 | 14 | 1.2893 | | | | | / | / |
| 6 | V2-R06-r6 | RWRIIAR | 7 | 0.8322 | 15 | 1.4567 | | | | | / | / |
| 7 | V2-R06-r6 | RWRIIAR | 7 | 0.8322 | 15 | 1.4567 | | | | | / | / |
| 8 | V2-R06-r6 | RWRIIAR | 7 | 0.8322 | 15 | 1.4567 | | | | | / | / |
| 9 | V2-R06-r6 | RWRIIAR | 7 | 0.8322 | 15 | 1.4567 | | | | | / | / |
| 10 | V2-R06-r6 | RWRIIAR | 7 | 0.8322 | 15 | 1.4567 | | | | | / | / |

[a] E-value against the AMPSphere database. "/" indicates no matches.
[b] E-value against the DRAMP database. "/" indicates no matches.
Continued on next page.

**Table S5:** Miniaturized sequences and scores during ten rounds of deletion by the deletion-only AM-PainterV2 (Continued).

| Round | Name | Sequence | Length | HyperAMP | #Del | \|△HyperAMP\| /#Del × 100 | MIC (μM) | | | | E-value[a] | E-value[b] |
|---|---|---|---|---|---|---|---|---|---|---|---|---|
| | | | | | | | Ec | Ab | Sa | Bs | | |
| 1 | V2-R07-r1 | LKIAKLNYFLIKGLLKKK | 18 | 0.8869 | 5 | 3.2742 | | | | | / | 5.1 |
| 2 | V2-R07-r2 | LKIAKLNLIKLLK | 13 | 0.8640 | 10 | 1.8659 | | | | | / | 4.6 |
| 3 | V2-R07-r2 | LKIAKLNLIKLLK | 13 | 0.8640 | 10 | 1.8659 | | | | | / | 4.6 |
| 4 | V2-R07-r2 | LKIAKLNLIKLLK | 13 | 0.8640 | 10 | 1.8660 | | | | | / | 4.6 |
| 5 | V2-R07-r2 | LKIAKLNLIKLLK | 13 | 0.8640 | 10 | 1.8660 | | | | | / | 4.6 |
| 6 | V2-R07-r2 | LKIAKLNLIKLLK | 13 | 0.8640 | 10 | 1.8660 | | | | | / | 4.6 |
| 7 | **V2-R07-D** | LKIAKLLIKLL | 11 | 0.8795 | 12 | **1.4258** | 128 | 128 | 16 | 32 | 4.4 | 0.8 |
| 8 | V2-R07-D | LKIAKLLIKLL | 11 | 0.8795 | 12 | 1.4258 | 128 | 128 | 16 | 32 | 4.4 | 0.8 |
| 9 | V2-R07-D | LKIAKLLIKLL | 11 | 0.8795 | 12 | 1.4258 | 128 | 128 | 16 | 32 | 4.4 | 0.8 |
| 10 | V2-R07-D | LKIAKLLIKLL | 11 | 0.8795 | 12 | 1.4258 | 128 | 128 | 16 | 32 | 4.4 | 0.8 |
| 1 | V2-R08-r1 | IPGTKWALKLAKKHMKLMKLIR | 22 | 1.0272 | 1 | 2.0852 | | | | | / | / |
| 2 | V2-R08-r2 | ITKWALKLAKKHMKLKLIR | 19 | 1.0025 | 4 | 1.1394 | | | | | / | / |
| 3 | V2-R08-r3 | ITKWALKLAKLKLIR | 15 | 1.0158 | 8 | 0.4037 | | | | | / | / |
| 4 | **V2-R08-D** | IKWALKLALKLIR | 13 | 1.0216 | 10 | **0.2650** | 32 | 64 | 4 | 8 | / | / |
| 5 | V2-R08-r5 | IKWALKLALLIR | 12 | 0.9414 | 11 | 0.9700 | | | | | / | / |
| 6 | V2-R08-r5 | IKWALKLALLIR | 12 | 0.9414 | 11 | 0.9700 | | | | | / | / |
| 7 | V2-R08-r5 | IKWALKLALLIR | 12 | 0.9414 | 11 | 0.9700 | | | | | / | / |
| 8 | V2-R08-r5 | IKWALKLALLIR | 12 | 0.9414 | 11 | 0.9700 | | | | | / | / |
| 9 | V2-R08-r9 | KWALKLALLIR | 11 | 0.9895 | 12 | 0.4883 | >128 | >128 | 64 | >128 | / | / |
| 10 | V2-R08-r10 | KALKLALLIR | 10 | 0.9356 | 13 | 0.8646 | | | | | / | / |
| 1 | V2-R09-r1 | IWNTIKRLPIWPWGTLRWPRL | 21 | 1.0096 | 1 | 2.8704 | | | | | / | 3.0 |
| 2 | V2-R09-r2 | IWNTIKRLIWPWGLRWRL | 18 | 0.9777 | 4 | 1.5151 | | | | | / | 7.0 |
| 3 | V2-R09-r3 | IWNTIKRLIWPWLRWRL | 17 | 1.0135 | 5 | 0.4947 | | | | | / | / |
| 4 | V2-R09-r3 | IWNTIKRLIWPWLRWRL | 17 | 1.0135 | 5 | 0.4940 | | | | | / | / |
| 5 | V2-R09-r5 | IWNTIKRLIWWLRWRL | 16 | 0.9771 | 6 | 1.0183 | | | | | / | / |
| 6 | V2-R09-r6 | INTIKRLIWWLRWRL | 15 | 0.9750 | 7 | 0.9043 | | | | | / | / |
| 7 | V2-R09-r6 | INTIKRLIWWLRWRL | 15 | 0.9750 | 7 | 0.9043 | | | | | / | / |
| 8 | **V2-R09-D** | NIKRLIWWLRWRL | 13 | 1.0256 | 9 | **0.1411** | 16 | 4 | 2 | 1 | / | / |
| 9 | V2-R09-r9 | NKRLIWWLRWRL | 12 | 0.9341 | 10 | 1.0420 | | | | | / | / |
| 10 | V2-R09-r9 | NKRLIWWLRWRL | 12 | 0.9341 | 10 | 1.0420 | | | | | / | / |
| 1 | V2-R10-r1 | GLLRWHWWKVHHLLKWIM | 18 | 0.9306 | 4 | 2.6184 | | | | | / | / |
| 2 | V2-R10-r2 | LLRWHWWKVHLLKWI | 15 | 0.9015 | 7 | 1.9120 | | | | | / | / |
| 3 | V2-R10-r3 | LLRWWKVHLLKWI | 13 | 0.9403 | 9 | 1.0556 | 8 | 8 | 2 | 2 | / | 6.7 |
| 4 | **V2-R10-D** | LLRWWKHLLKWI | 12 | 0.9328 | 10 | **1.0250** | 4 | 2 | 2 | 2 | / | / |
| 5 | V2-R10-r5 | LLWKHLLKWI | 10 | 0.8347 | 12 | 1.6725 | | | | | 9.6 | / |
| 6 | V2-R10-r5 | LLWKHLLKWI | 10 | 0.8347 | 12 | 1.6725 | | | | | 9.6 | / |
| 7 | V2-R10-r5 | LLWKHLLKWI | 10 | 0.8347 | 12 | 1.6725 | | | | | 9.6 | / |
| 8 | V2-R10-r8 | LLWKHLLKI | 9 | 0.8108 | 13 | 1.7277 | | | | | 4.7 | 3.2 |
| 9 | V2-R10-r8 | LLWKHLLKI | 9 | 0.8108 | 13 | 1.7277 | | | | | 4.7 | 3.2 |
| 10 | V2-R10-r8 | LLWKHLLKI | 9 | 0.8108 | 13 | 1.7277 | | | | | 4.7 | 3.2 |
| 1 | V1-R04-r1 | RGWKKRRIKRFMRGLRGFIKAA | 22 | 1.1732 | 4 | 0.7155 | | | | | / | 7.1 |
| 2 | **V1-R04-D** | RWKKRIKRFMRGLRFIKAA | 19 | 1.1107 | 7 | **0.4839** | 1 | 2 | 1 | 2 | / | 4.4 |
| 3 | V1-R04-r3 | RKKRIRMRLRFIKA | 14 | 0.9956 | 12 | 1.2419 | | | | | / | / |
| 4 | V1-R04-r4 | RKKRIRLRIKA | 11 | 0.9882 | 15 | 1.0427 | | | | | / | 0.7 |
| 5 | V1-R04-r4 | RKKRIRLRIKA | 11 | 0.9882 | 15 | 1.0427 | | | | | / | 0.7 |
| 6 | V1-R04-r6 | RKRIRLRIKA | 10 | 1.0226 | 16 | 0.7625 | >128 | >128 | 128 | >128 | / | 0.6 |
| 7 | V1-R04-r6 | RKRIRLRIKA | 10 | 1.0226 | 16 | 0.7627 | >128 | >128 | 128 | >128 | / | 0.6 |
| 8 | V1-R04-r6 | RKRIRLRIKA | 10 | 1.0226 | 16 | 0.7625 | >128 | >128 | 128 | >128 | / | 0.6 |
| 9 | V1-R04-r6 | RKRIRLRIKA | 10 | 1.0226 | 16 | 0.7625 | >128 | >128 | 128 | >128 | / | 0.6 |
| 10 | V1-R04-r6 | RKRIRLRIKA | 10 | 1.0226 | 16 | 0.7625 | >128 | >128 | 128 | >128 | / | 0.6 |

[a] E-value against the AMPSphere database. "/" indicates no matches.
[b] E-value against the DRAMP database. "/" indicates no matches.

### Supplementary Figures

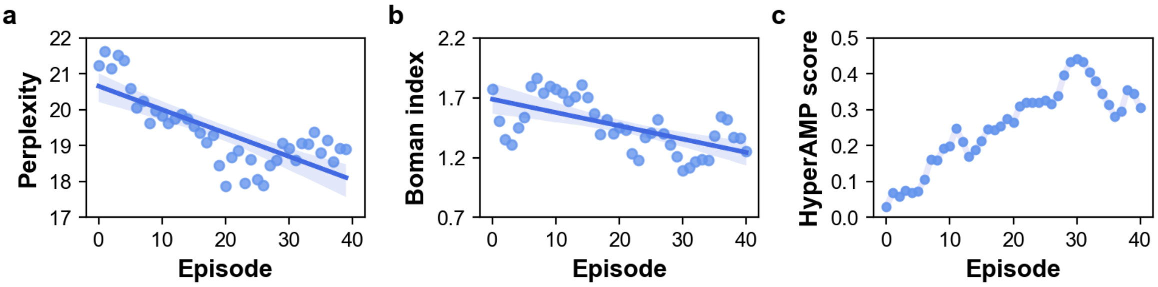


**Figure S1:** Training process of AMPainterV2. **a**. Perplexity decreases gradually during training for the first 30 episodes. **b**. Boman index of sequences declines. **c**. The HyperAMP score increases initially and then declines slightly, with the highest score in episode 30.

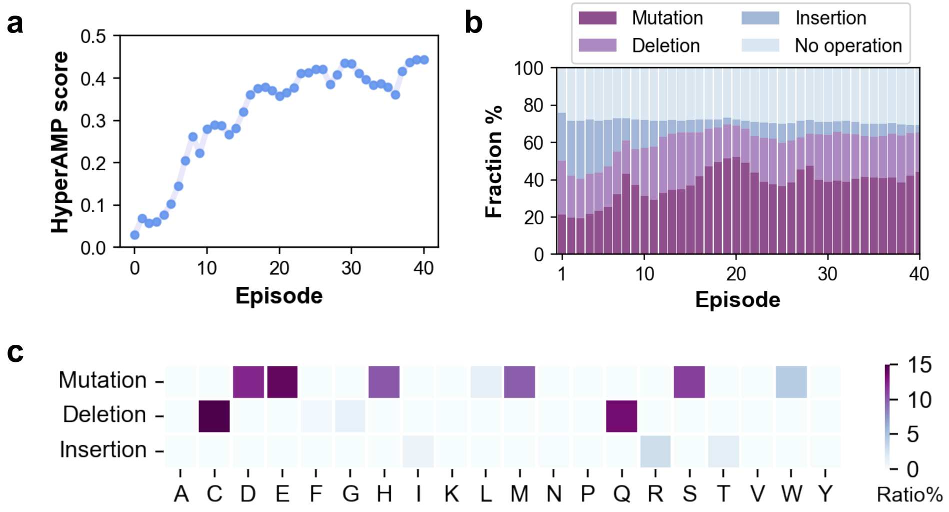


**Figure S2:** AMPainterV2 without adding noise in the MoE policy. **a**. HyperAMP score during the training stage. **b**. Expert preference. **c**. Assigned amino acid preference of each expert at episode 40.

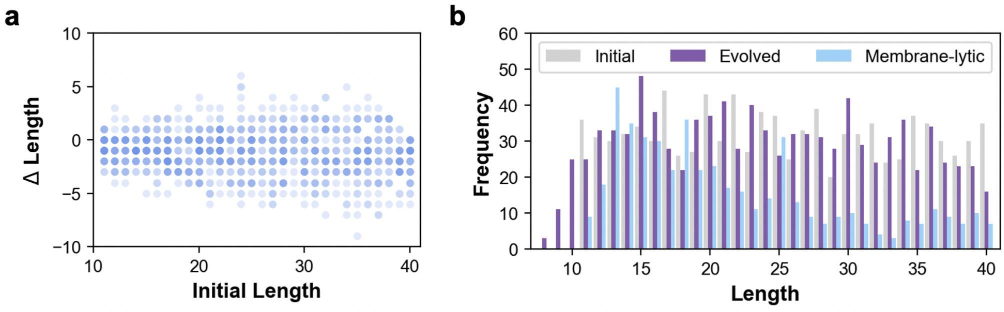


**Figure S3:** Sequence length distributions. **a**. ΔLength of evolved sequences versus the length of their initial sequences. **b**. Length distribution of initial random sequences, evolved sequences at episode 30, and the membrane-lytic AMPs.

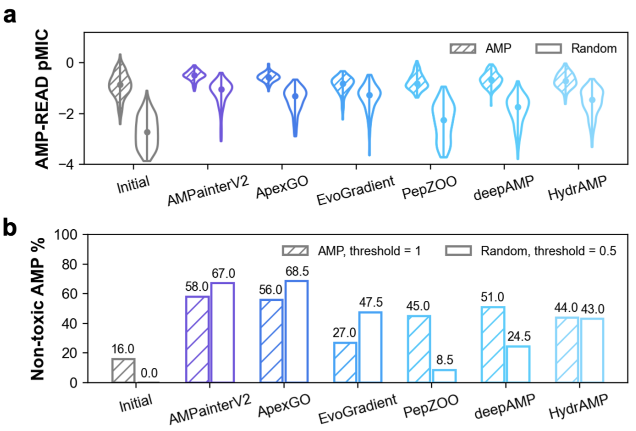


**c**

| Method | Diversity | | Similarity (%) | |
|---|---|---|---|---|
| | AMP | Random | AMP | Random |
| Initial | 17.036 | 17.482 | 48.58 | 41.30 |
| AMPainterV2 | **17.099** | **17.842** | **44.14** | **45.43** |
| ApexGO | 16.573 | 16.772 | 48.09 | 43.79 |
| EvoGradient | 15.642 | 17.288 | 48.51 | 42.63 |
| PepZOO | 15.698 | 16.009 | 48.42 | 42.52 |
| deepAMP | 15.994 | 16.500 | 49.20 | 44.08 |
| HydrAMP | 16.431 | 16.794 | 47.98 | 43.34 |

**Figure S4:** Additional comparison results. **a**. AMP-READ pMIC scores of the initial and evolved sequences. **b**. Ratio of non-toxic AMPs in initial and evolved sequences. Toxicity was evaluated by ToxinPred 3.0. **c**. Diversity and similarity metrics of initial and evolved sequences. The metrics of AMPainterV2 are shown in bold.

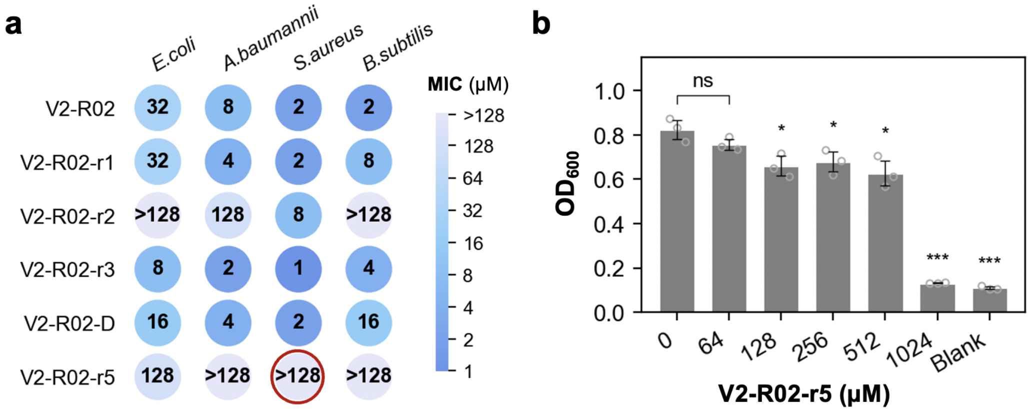


**Figure S5:** Shortened peptides of V2-R02 by deletion-only AMPainterV2. **a**. MICs against four bacterial strains. **b**. Absorbance at 600 nm of V2-R02-r5 in the MIC measurement against *S. aureus*, showing the MIC value of 1024 μM. V2-R02-r5 was dissolved in sterile water to exclude the influence of high-concentration DMSO. Unpaired t-tests were used for comparison with 0 μM. ns, no significant. *, $p < 0.05$. ***, $p < 0.001$.

**a**

| | | |
|---|---|---|
| V2-R04 | KSLYTWVKKIKHRIYKMWIWFGKHIF | 26 |
| V2-R04-r1 | K-L-TW-KKI-HRI-KMWIWF-KHIF | 20 |
| V2-R04-D | K-L-TW-KKI--RI-K-WI---KHIF | 16 |
| V2-R04-r3 | K-L-TW--KI--RI-K--I---K-I- | 12 |
| V2-R04-r5 | K-L-----KI--RI-K--I-----I- | 9 |
| V2-R04-r6 | K-L-----KI--RI-K--I------- | 8 |
| V2-R04-r8 | K-L-----KI--RI----I------- | 7 |

**b** **c**

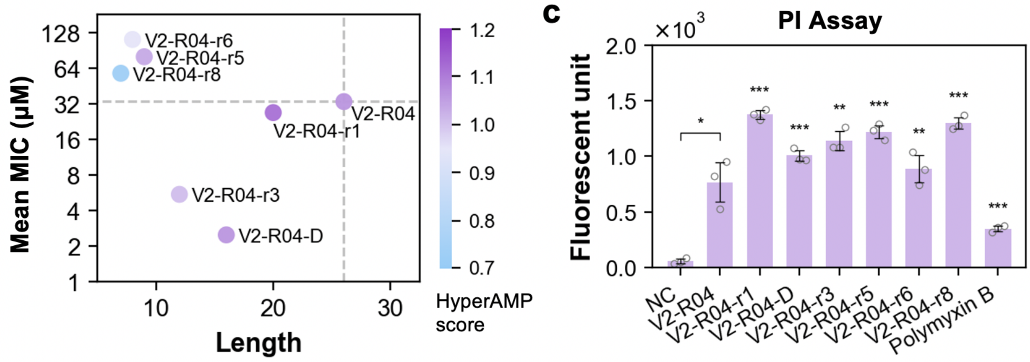


**Figure S6:** Shortened peptides of V2-R04 by deletion-only AMPainterV2. **a**. Names, sequences, and lengths of peptides. **b**. Mean MICs against four bacterial strains versus the lengths of these peptides. The color bar shows the HyperAMP score of each peptide. **c**. Membrane-lytic ability measured via the PI assay on *S. aureus*. Unpaired t-tests were used for comparison with the negative control (NC). *, $p < 0.05$. **, $p < 0.01$. ***, $p < 0.001$.

**a** **b**

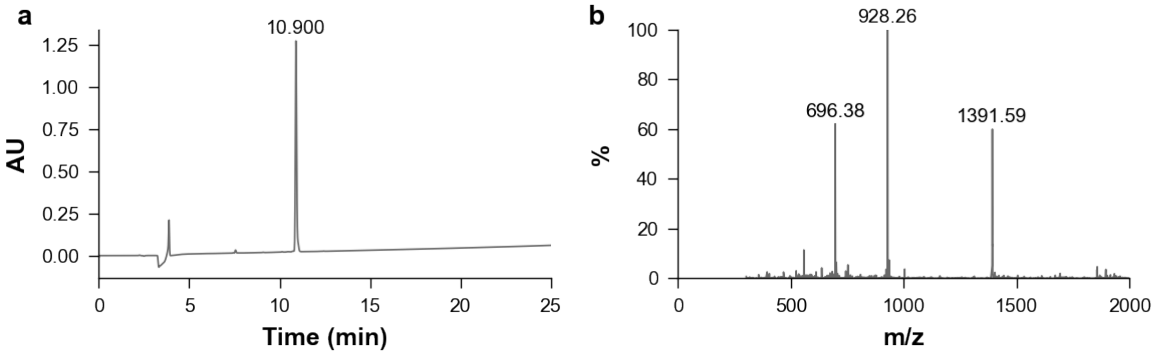


**Figure S7:** Validation of synthesized peptide V2-R01. **a**, HPLC. **b**, Mass spectrometry.

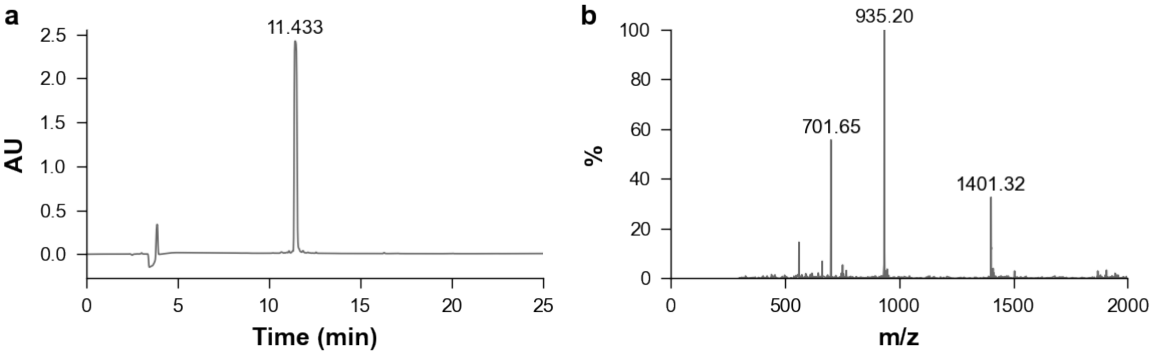


**Figure S8:** Validation of synthesized peptide V2-R02. **a**. HPLC. **b**. Mass spectrometry.

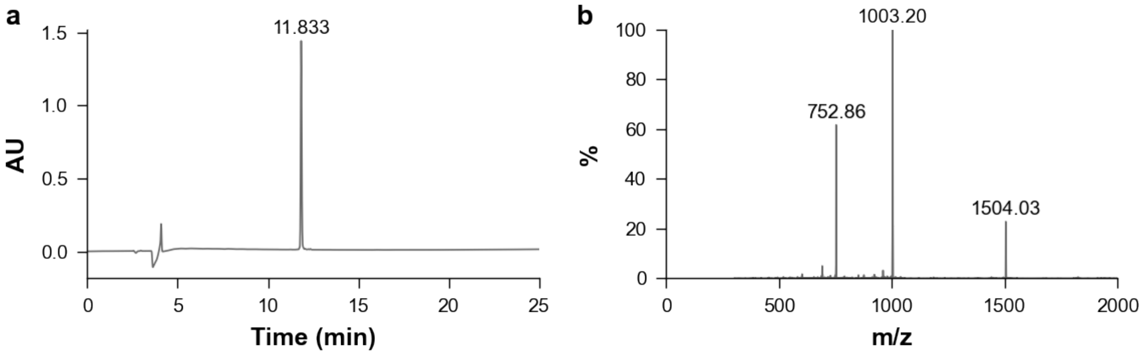


**Figure S9:** Validation of synthesized peptide V2-R03. **a**. HPLC. **b**. Mass spectrometry.

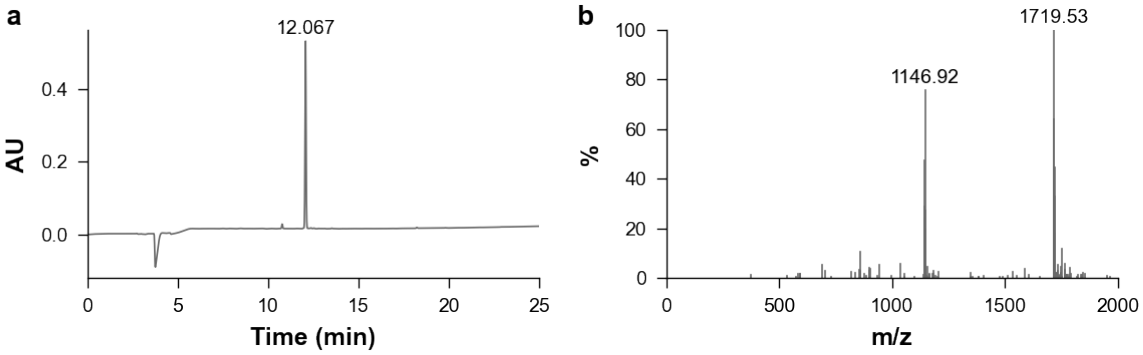


**Figure S10:** Validation of synthesized peptide V2-R04. **a**. HPLC. **b**. Mass spectrometry.

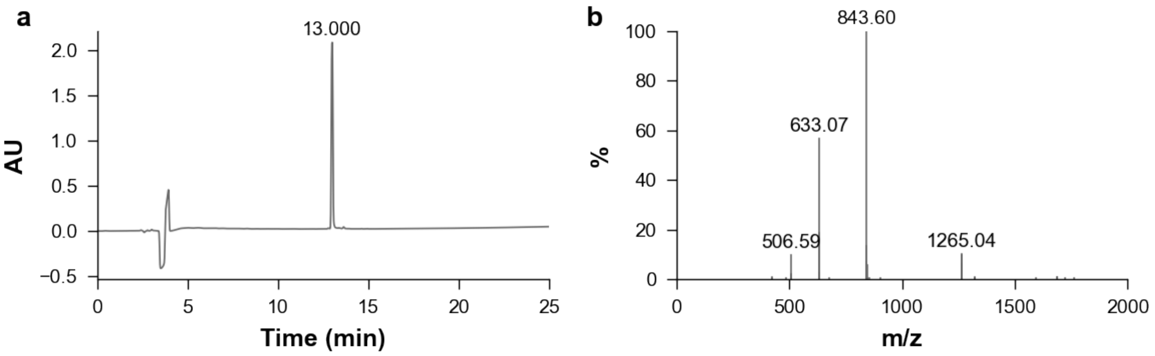


**Figure S11:** Validation of synthesized peptide V2-R05. **a**. HPLC. **b**. Mass spectrometry.

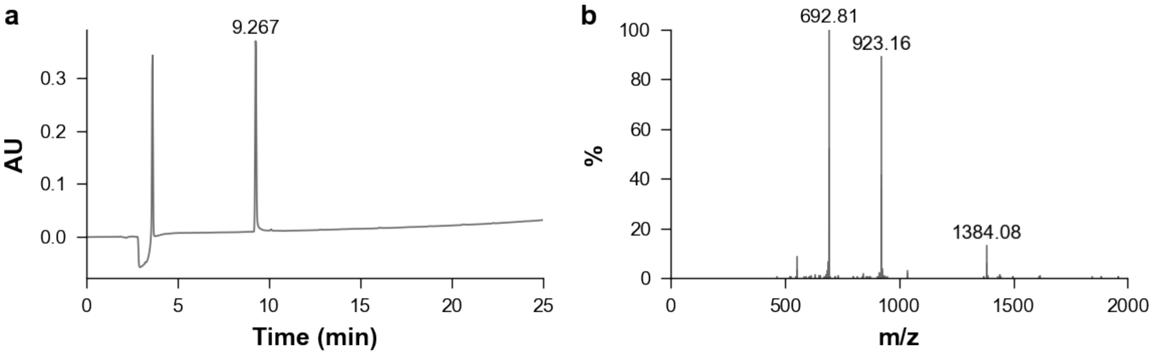


**Figure S12:** Validation of synthesized peptide V2-R06. **a**. HPLC. **b**. Mass spectrometry.

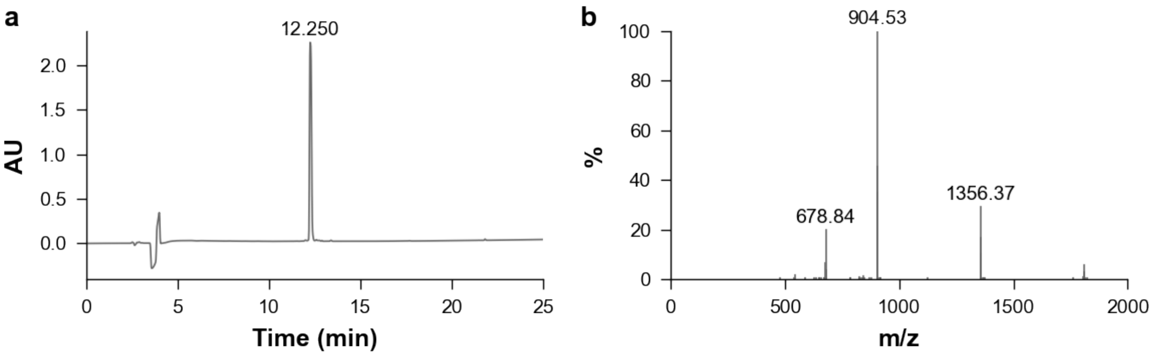


**Figure S13:** Validation of synthesized peptide V2-R07. **a**. HPLC. **b**. Mass spectrometry.

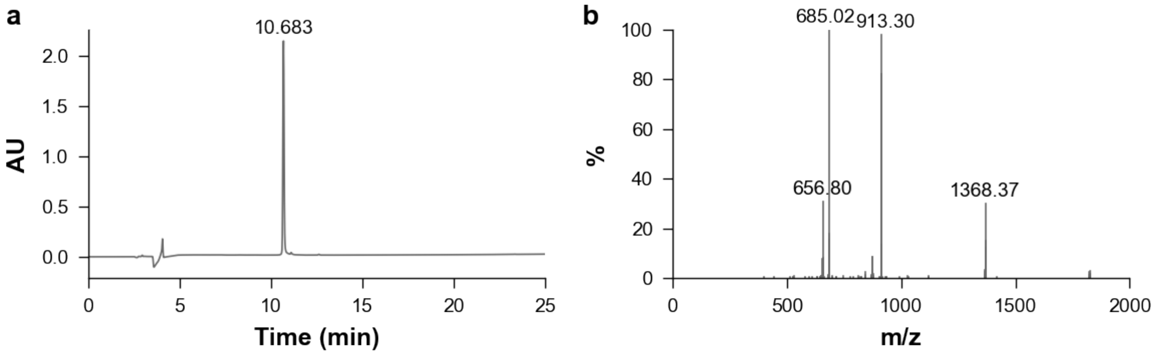


**Figure S14:** Validation of synthesized peptide V2-R08. **a**. HPLC. **b**. Mass spectrometry.

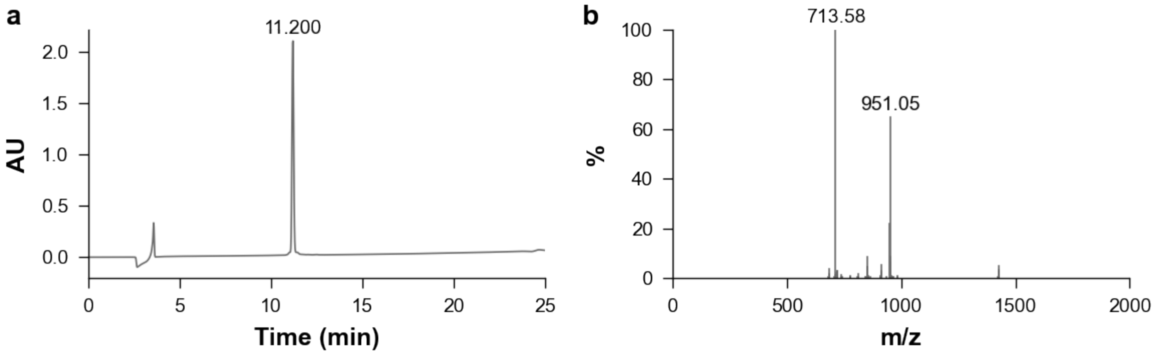


**Figure S15:** Validation of synthesized peptide V2-R09. **a**. HPLC. **b**. Mass spectrometry.

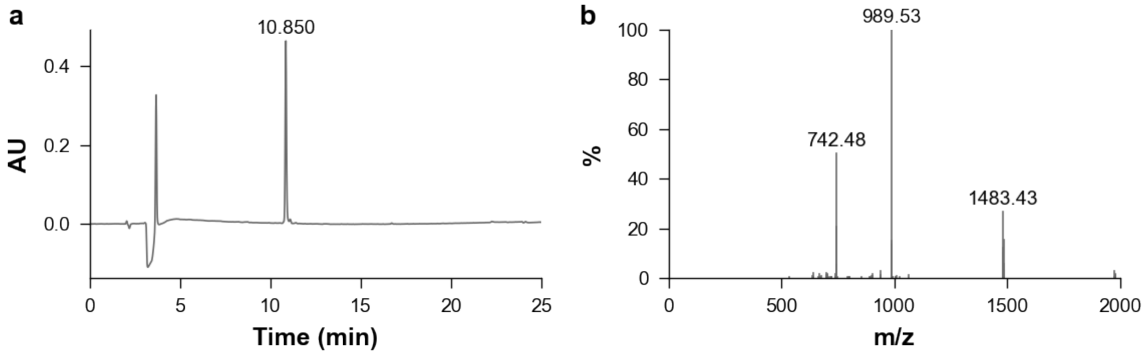


**Figure S16:** Validation of synthesized peptide V2-R10. **a**. HPLC. **b**. Mass spectrometry.

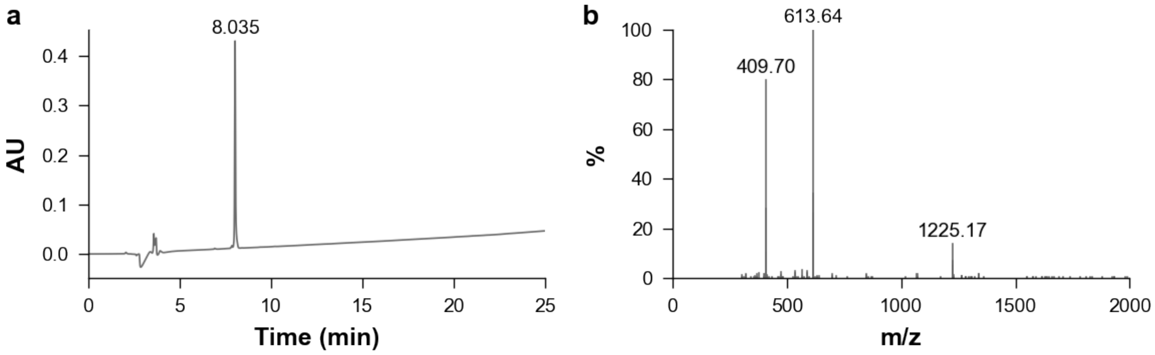


**Figure S17:** Validation of synthesized peptide V2-R01-D. **a**. HPLC. **b**. Mass spectrometry.

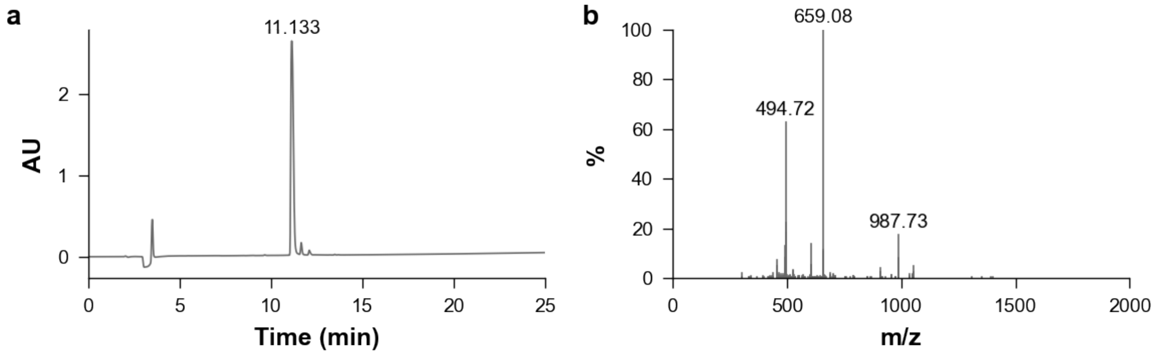


**Figure S18:** Validation of synthesized peptide V2-R01-r1. **a**. HPLC. **b**. Mass spectrometry.

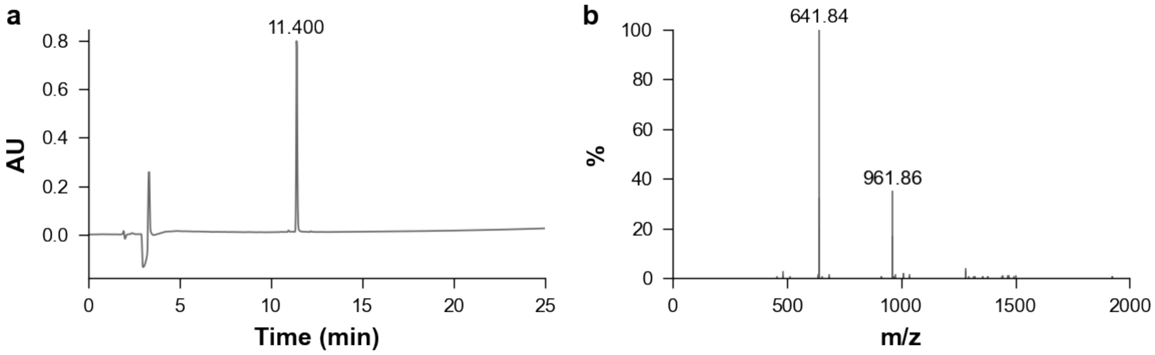


**Figure S19:** Validation of synthesized peptide V2-R02-D. **a**. HPLC. **b**. Mass spectrometry.

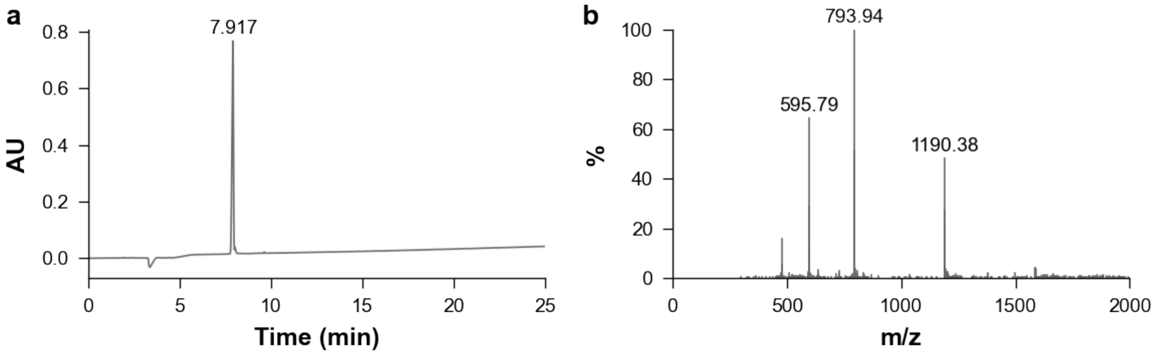


**Figure S20:** Validation of synthesized peptide V2-R02-r1. **a**. HPLC. **b**. Mass spectrometry.

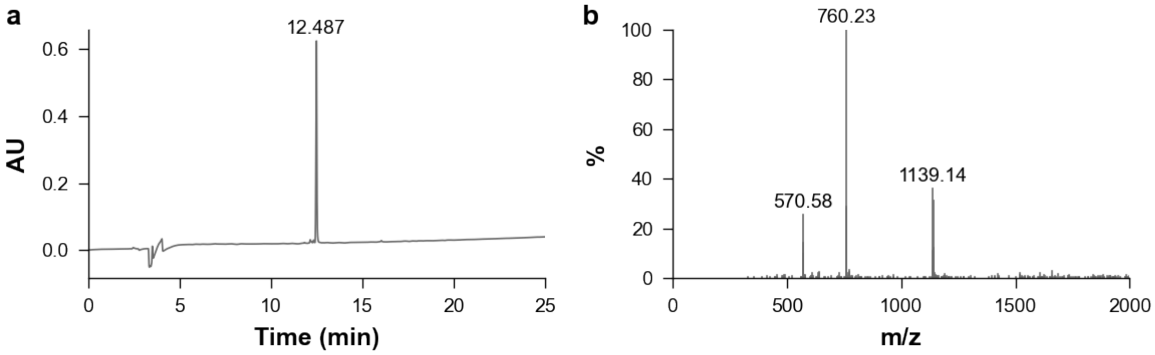


**Figure S21:** Validation of synthesized peptide V2-R02-r2. **a**. HPLC. **b**. Mass spectrometry.

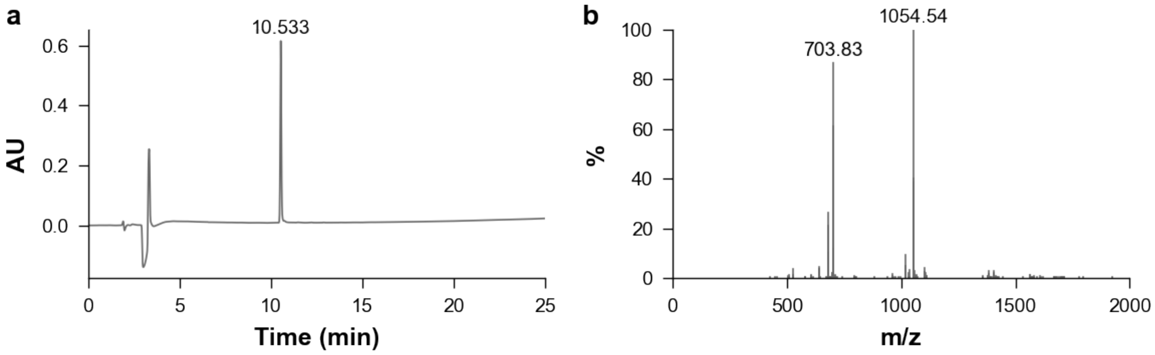


**Figure S22:** Validation of synthesized peptide V2-R02-r3. **a**. HPLC. **b**. Mass spectrometry.

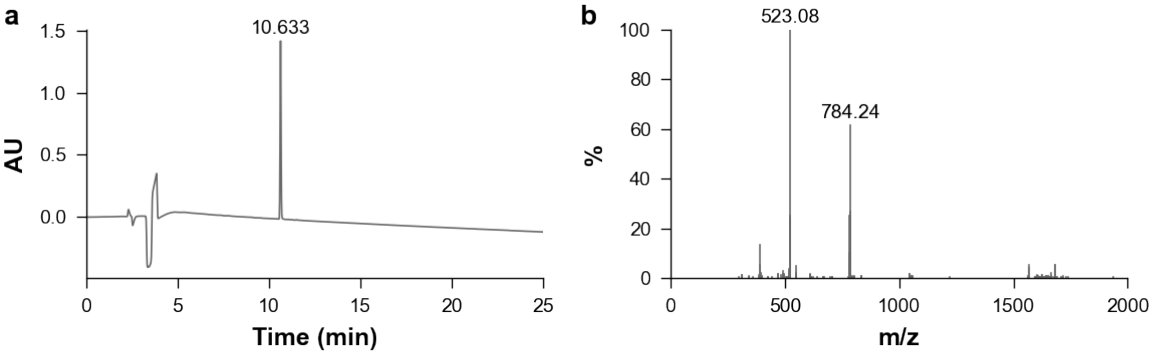


**Figure S23:** Validation of synthesized peptide V2-R02-r5. **a**. HPLC. **b**. Mass spectrometry.

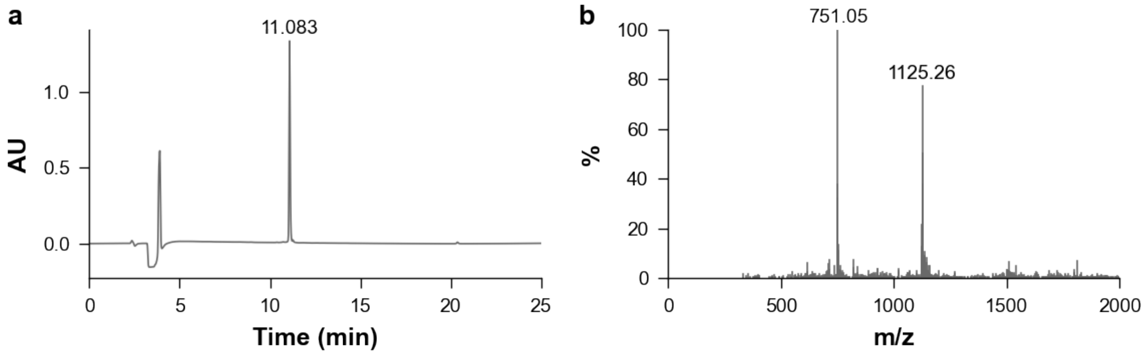


**Figure S24:** Validation of synthesized peptide V2-R03-D. **a**. HPLC. **b**. Mass spectrometry.

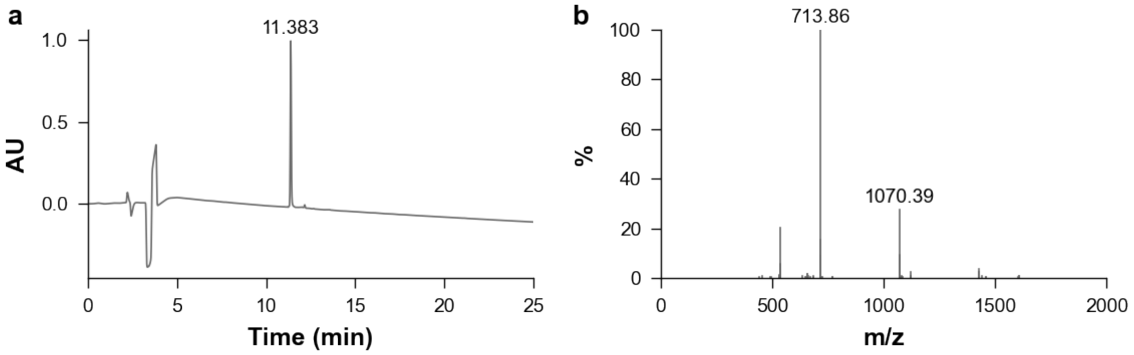


**Figure S25:** Validation of synthesized peptide V2-R04-D. **a**. HPLC. **b**. Mass spectrometry.

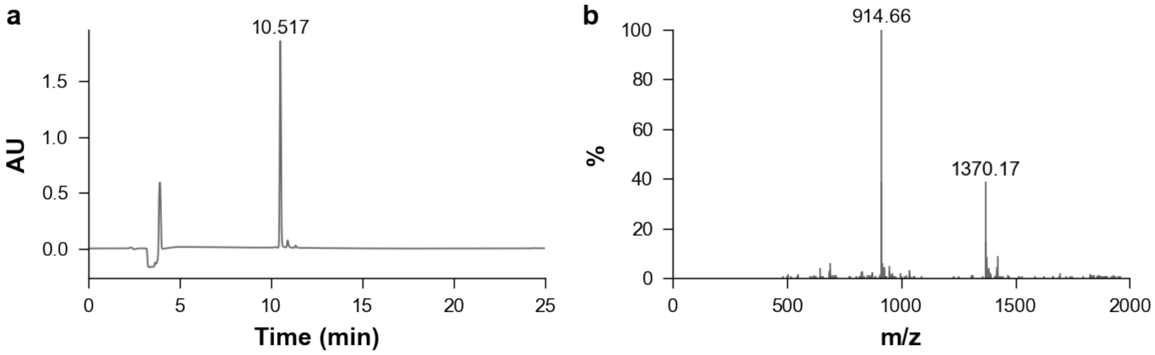


**Figure S26:** Validation of synthesized peptide V2-R04-r1. **a**. HPLC. **b**. Mass spectrometry.

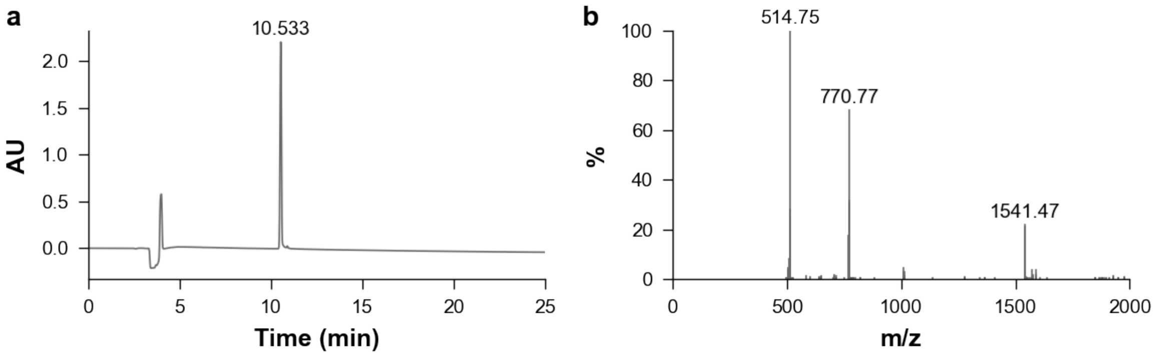


**Figure S27:** Validation of synthesized peptide V2-R04-r3. **a**. HPLC. **b**. Mass spectrometry.

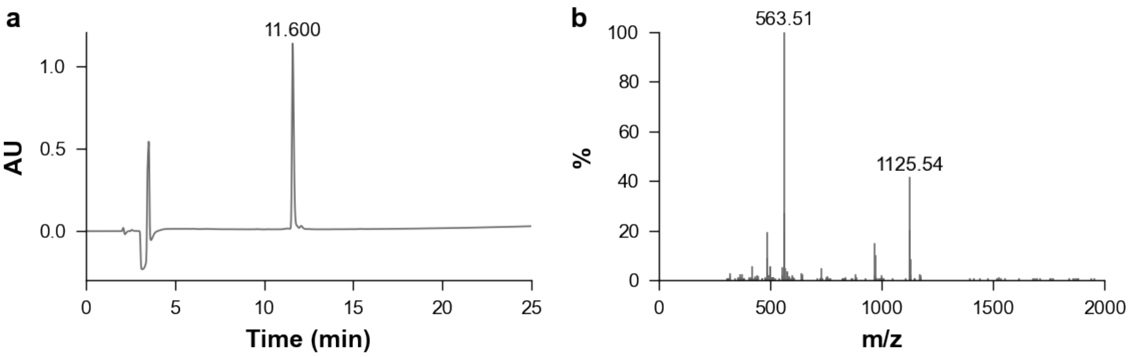


**Figure S28:** Validation of synthesized peptide V2-R04-r4. **a**. HPLC. **b**. Mass spectrometry.

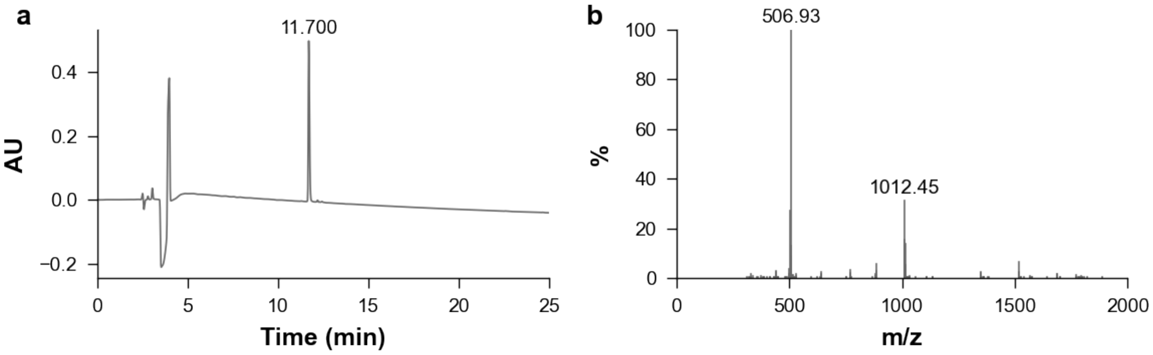


**Figure S29:** Validation of synthesized peptide V2-R04-r6. **a**. HPLC. **b**. Mass spectrometry.

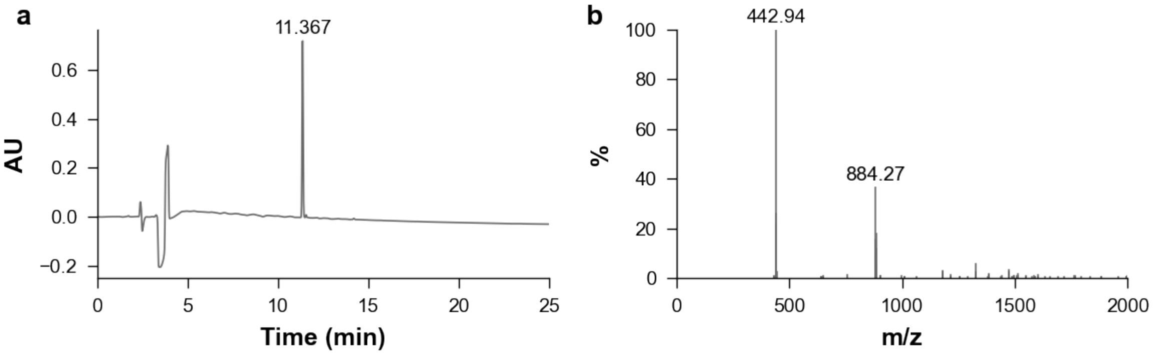


**Figure S30:** Validation of synthesized peptide V2-R04-r8. **a**. HPLC. **b**. Mass spectrometry.

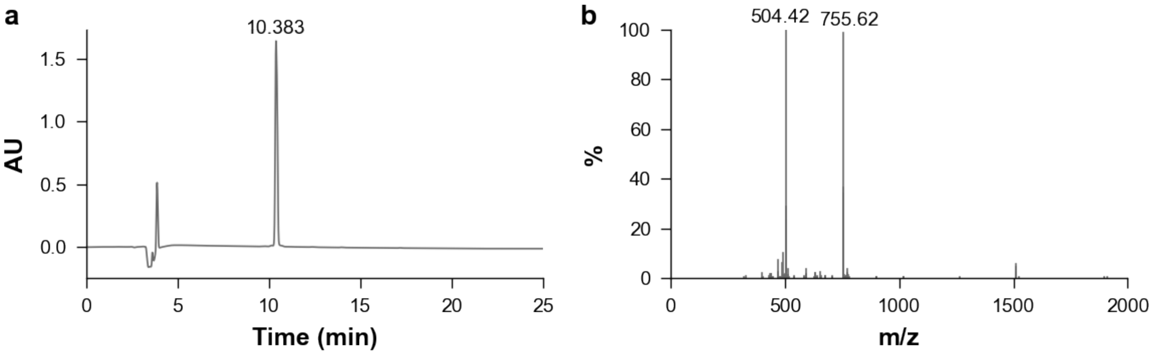


**Figure S31:** Validation of synthesized peptide V2-R05-D. **a**. HPLC. **b**. Mass spectrometry.

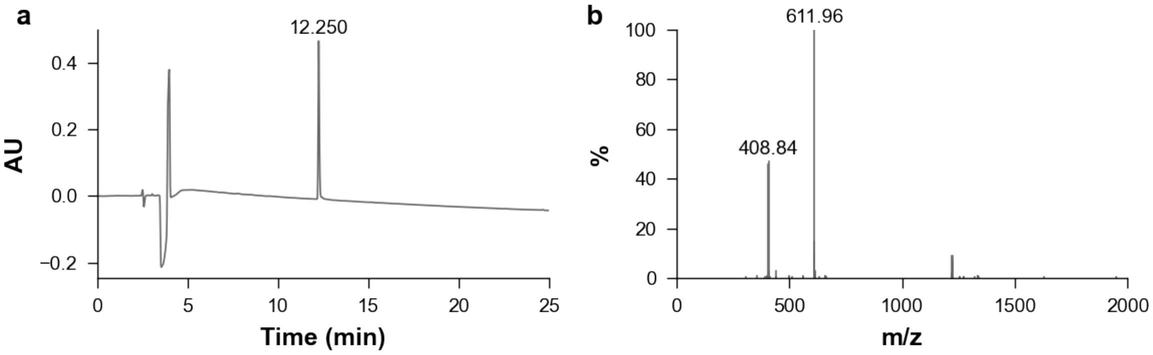


**Figure S32:** Validation of synthesized peptide V2-R05-r10. **a**. HPLC. **b**. Mass spectrometry.

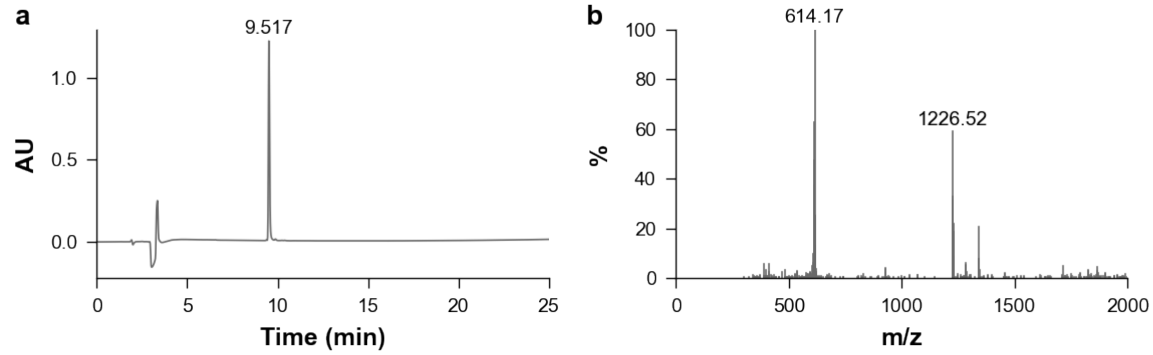


**Figure S33:** Validation of synthesized peptide V2-R06-D. **a**. HPLC. **b**. Mass spectrometry.

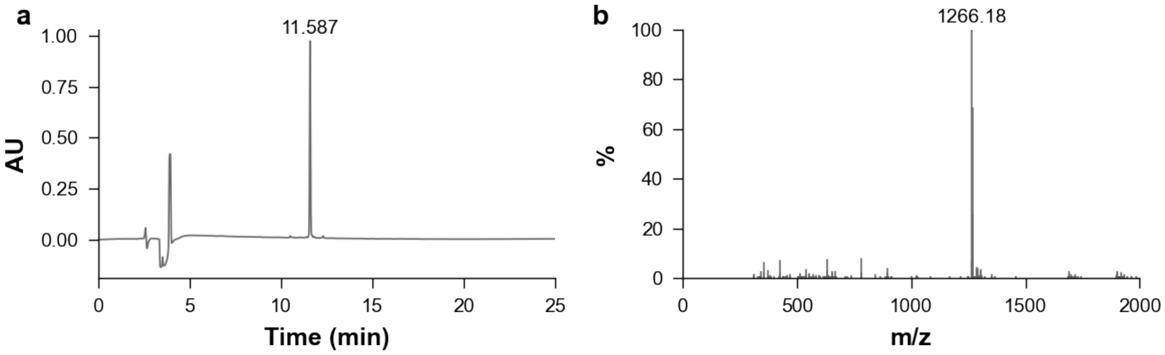


**Figure S34:** Validation of synthesized peptide V2-R07-D. **a**. HPLC. **b**. Mass spectrometry.

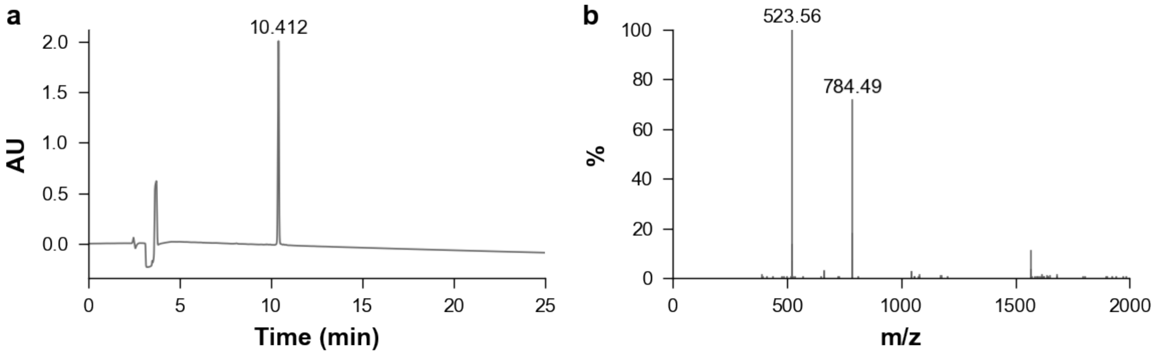


**Figure S35:** Validation of synthesized peptide V2-R08-D. **a**. HPLC. **b**. Mass spectrometry.

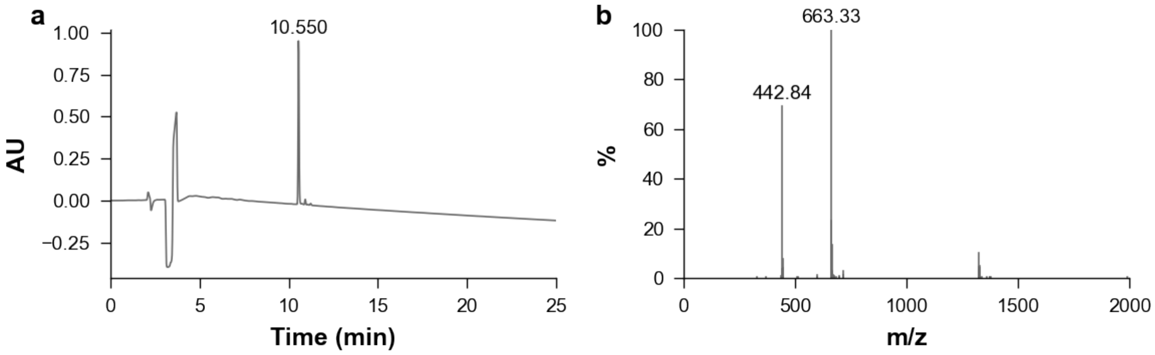


**Figure S36:** Validation of synthesized peptide V2-R08-r9. **a**. HPLC. **b**. Mass spectrometry.

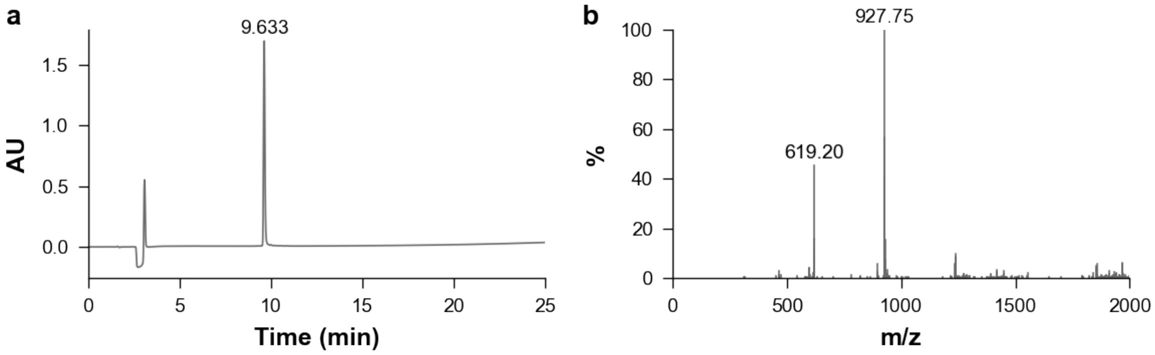


**Figure S37:** Validation of synthesized peptide V2-R09-D. **a**. HPLC. **b**. Mass spectrometry.

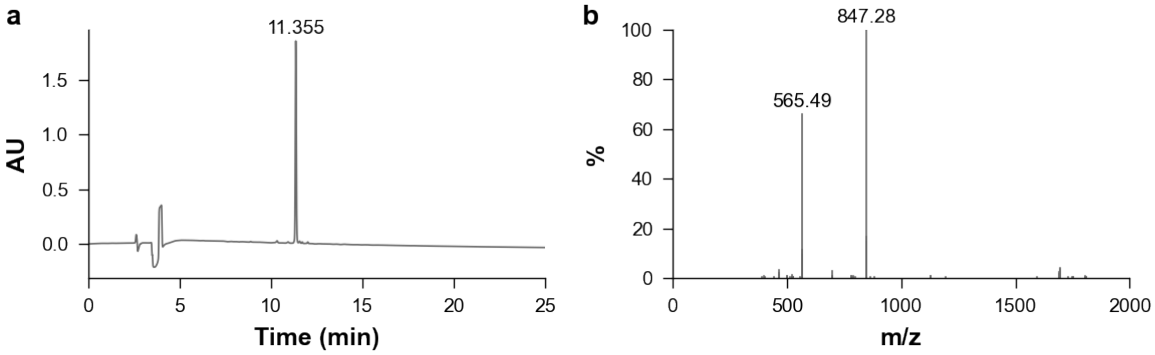


**Figure S38:** Validation of synthesized peptide V2-R10-D. **a**. HPLC. **b**. Mass spectrometry.

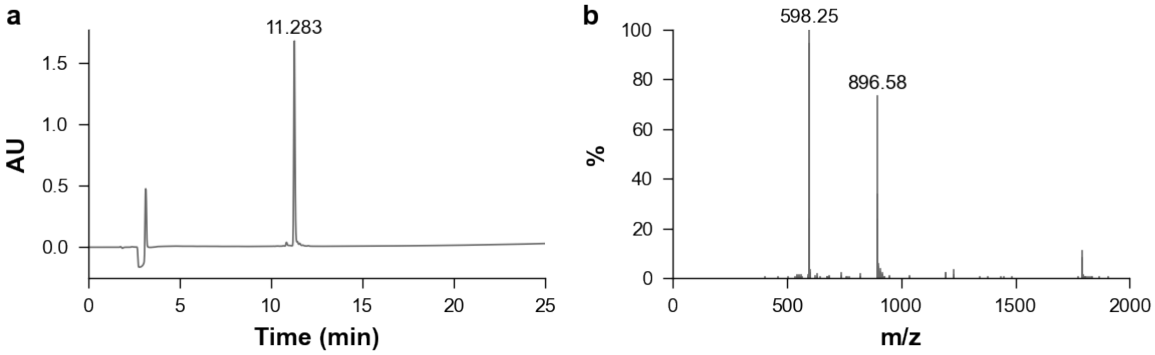


**Figure S39:** Validation of synthesized peptide V2-R10-r3. **a**. HPLC. **b**. Mass spectrometry.

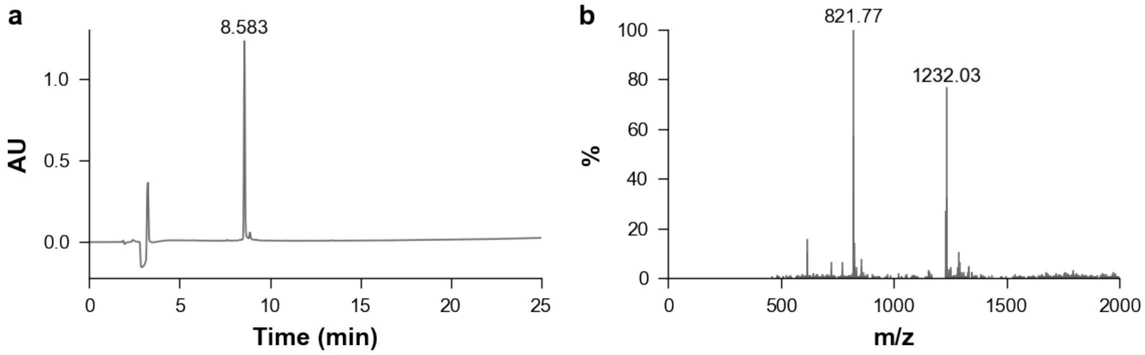


**Figure S40:** Validation of synthesized peptide V1-R04-D. **a**. HPLC. **b**. Mass spectrometry.

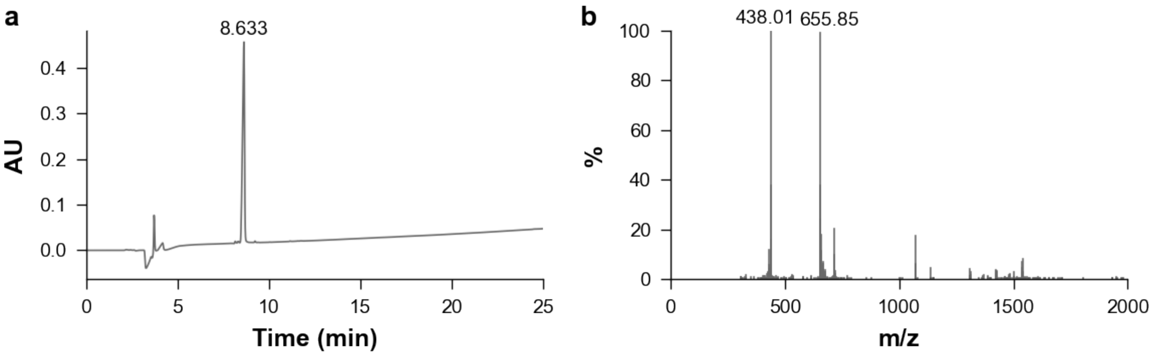


**Figure S41:** Validation of synthesized peptide V1-R04-r6. **a**. HPLC. **b**. Mass spectrometry.
